# Strength of species interactions determines biodiversity and stability in microbial communities

**DOI:** 10.1101/671008

**Authors:** Christoph Ratzke, Julien Barrere, Jeff Gore

**Author notes:** equal contribution (Christoph Ratzke, Julien Barrere). correspondence should be sent to or.

## Abstract

Organisms – especially microbes – tend to live in complex communities. While some of these ecosystems are very bio-diverse, others aren’t^1–3^, and while some are very stable over time others undergo strong temporal fluctuations^4,5^. Despite a long history of research and a plethora of data it is not fully understood what sets biodiversity and stability of ecosystems^6,7^. Theory as well as experiments suggest a connection between species interaction, biodiversity, and stability of ecosystems^8–13^, where an increase of ecosystem stability with biodiversity could be observed in several cases^7,9,14^. However, what causes these connections remains unclear. Here we show in microbial ecosystems in the lab that the concentrations of available nutrients can set the strength of interactions between bacteria. At high nutrient concentrations, extensive microbial growth leads to strong chemical modifications of the environment, causing more negative interactions between species. These stronger interactions exclude more species from the community – resulting in a loss of biodiversity. At the same time, these stronger interactions also decrease the stability of the microbial communities, providing a mechanistic link between species interaction, biodiversity and stability.

## Main

Interactions between microbes are basic building blocks of microbial ecosystems^15–17^. They strongly influence who is present or absent in the community and therefore set the overall composition, stability and biodiversity of microbial ecosystems (Fig. 1A). Thus, it should be possible to understand microbial communities from bacterial interactions using a bottom-up approach^18^. However, how all these microbial interactions work together remains unresolved, which raises the question of whether we can gain insight into complex communities from studying simple microbial interactions at all. We show in the following that we could indeed transfer basic properties of simple interactions to large microbial assemblages and this way mechanistically understand what determines biodiversity and stability in several complex microbial communities.

**Figure 1:**
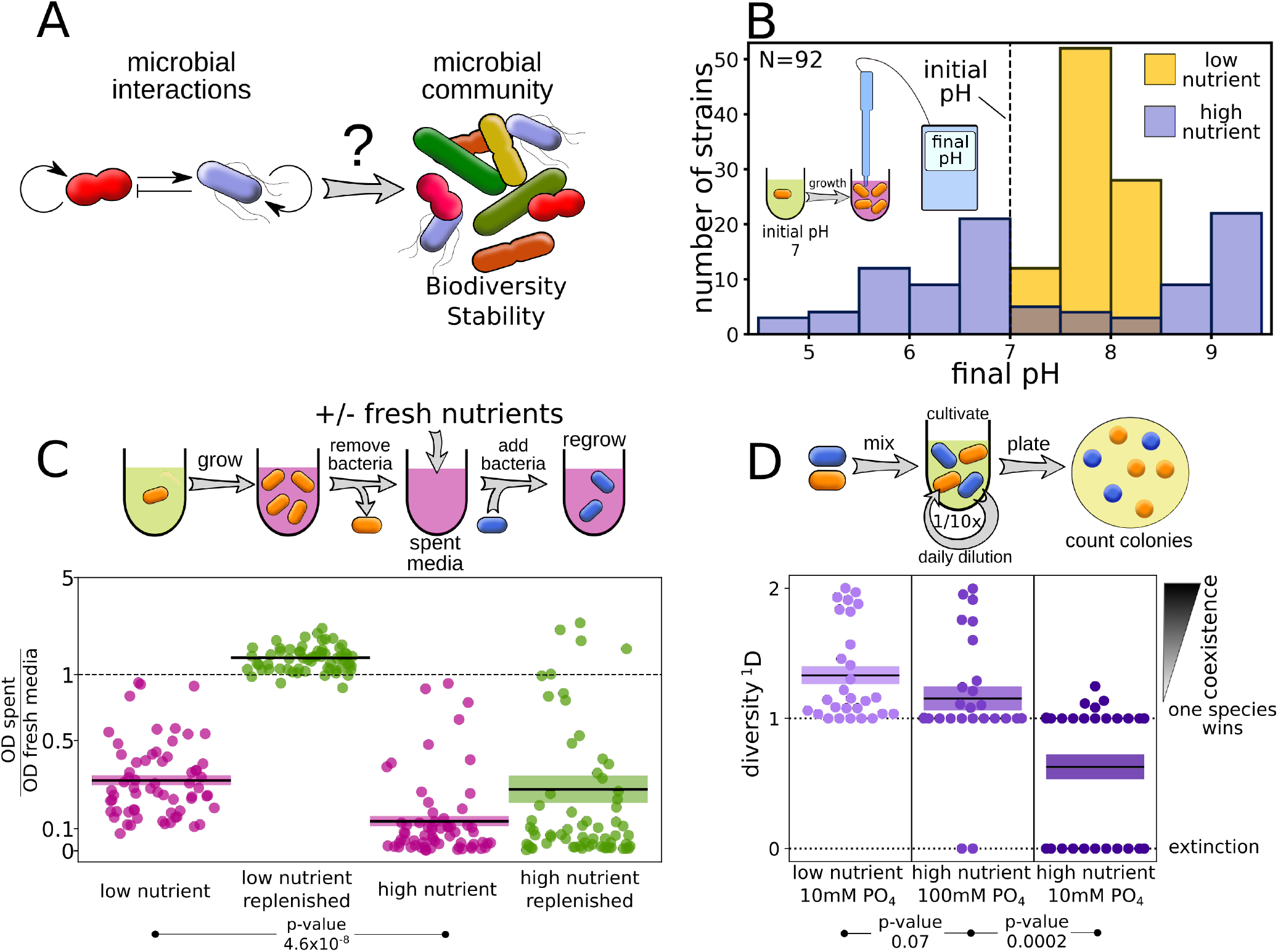
Higher nutrient concentrations lead to stronger negative interactions between microbes. (A) Can we understand biodiversity and stability of complex microbial communities from simple bacterial interactions? (B) Bacteria change the environmental pH stronger at higher nutrient concentrations. (C) Spent media of different bacteria were used either directly (purple) or after replenishing the resources (green) to re-grow the bacteria. All 64 pairs are shown separately in Supplementary Fig. 4. The plot shows relative growth for every interaction pair as scatter plot and the means +/− SEM as boxes. (D) Accordingly high nutrient concentrations decrease coexistence between interacting pairs. Low nutrient means 0.1% yeast extract, 0.1% soytone. High nutrient is the same medium with additional 1% glucose and 0.8% urea. All 28 co-culture outcomes are shown as swarm plot and the means +/− SEM as boxes. For more detailed information see the methods section. p-values were calculated with one-sided t-test. The diversity is calculated with 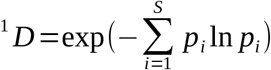, where p_i_ is the relative abundance of species i. If both species went extinct ^1^D was set to 0.

Microbes interact in many ways; they can compete for resources, inhibit each other by the production of antibiotics, or support each other via cross-feeding^15,19^. Most of these interactions are mediated through the environment: bacteria chemically modify their surroundings, which directly influences them as well as other members of the community. We and others recently showed that interactions between microbes can be understood and even predicted by understanding how they modify and react to their environment^19–23^. The higher the nutrient concentrations the microbes have access to the stronger they can metabolize and hence the stronger they can modify the environment. Accordingly, we expect that higher nutrient concentrations lead to stronger interactions, which may have a strong impact on essential ecosystem properties, like biodiversity and stability^8,13^.

To explore this idea, we first studied how interaction strength is influenced by nutrient concentrations in the context of pairwise interactions. An important environmental parameter that all microbes influence and are influenced by is the pH. The pH is altered by the uptake and production of many different substances and therefore delivers an integral metric of how the bacteria change their environment. Since different bacteria reach maximum growth at different pH values (Supplementary Fig. 1), by changing the pH they can directly impact their own and others’ growth. We measured the change of the environmental pH by 92 soil bacteria (Supplementary Fig. 2B) in 0.1% yeast extract, 0.1% soytone with or without additional 1% glucose and 0.8% urea. We will refer to these two conditions as high and low nutrient concentrations respectively. When grown at low nutrient concentrations with an initial pH of 7, bacteria slightly shifted the pH of the media towards the alkaline, whereas at high nutrient concentrations they either strongly increased or decreased the pH (Fig. 1B). As expected, stronger buffering or intermediate nutrient concentrations lead to intermediate pH change (Supplementary Fig. 2).

To test if this stronger change of the environment at high nutrient concentrations also increases interaction strength we grew 8 different soil bacteria (Supplementary Fig. 3) at low and high nutrient concentrations then took their spent media and re-grew each of the species in the spent media of the others (Fig. 1C, left panel). Bacterial growth on spent media from low nutrient media usually lowered the growth but did not completely inhibit it. This growth effect could be attenuated by adding fresh nutrients to the spent media, showing that the growth inhibition was largely driven by resource competition. On the other hand, spent media from high nutrient concentrations led to even more pronounced negative interactions and repressed bacterial growth completely in many cases, although in 10 out of 64 cases a relative facilitation was instead observed (Supplementary Fig. 4). Unlike our observation for low nutrient conditions, this growth inhibition at high nutrient concentrations could not be overcome by the addition of fresh nutrients (Fig. 1C, right panel). Therefore, the negative interactions are mostly driven by the production of toxic metabolites and not by the competition for resources. Buffering the media removed a large fraction of the inhibitory effect of the supernatant, suggesting that pH was a major factor causing this toxicity (Supplementary Fig. 5). Overall, our bacteria produced a more harmful environment when grown at higher nutrient concentrations.

To determine the consequence of these environmental modifications on pairwise coexistence, we co-cultured all pairwise combinations of the 8 species in batch culture with daily dilution in both low and high nutrient condition (Fig. 1D). After 5 days, the composition of the cultures was assayed by plating the bacteria and counting the different colonies (see methods for details). At low nutrient concentrations, there was a high amount of coexistence in pairwise co-culture. For the same interaction partners at high nutrient concentrations we observed a striking loss of coexistence, where either one species out-competed the other or, in many cases, both went extinct by ecological suicide as we described recently^21^. Intermediate nutrient concentrations lead to intermediate loss of coexistence (Supplementary Fig. 6). Higher buffer concentrations prevented the loss of coexistence at high nutrient concentrations, showing once more that pH is a major driver of the species interactions (Fig. 1D, lower middle). A similar but weaker loss of coexistence at high nutrient concentrations was also observed when increasing the concentrations of complex nutrients (Supplementary Fig. 7). Therefore, an increase in nutrient concentrations led to an increase in interaction strength, resulting in a loss of coexistence.

To explore whether these dynamics play out in complex communities, we sampled several soil microbiotas: compost, soil from an indoor flowerpot and soil from a local backyard. Those samples were cultivated in low and high nutrient conditions as described above, with daily dilutions into fresh media (see methods for details). The composition of the communities was followed over time by taking samples every day and performing 16S rRNA amplicon sequencing (Fig. 2 and Supplementary Fig. 9).

**Figure 2:**
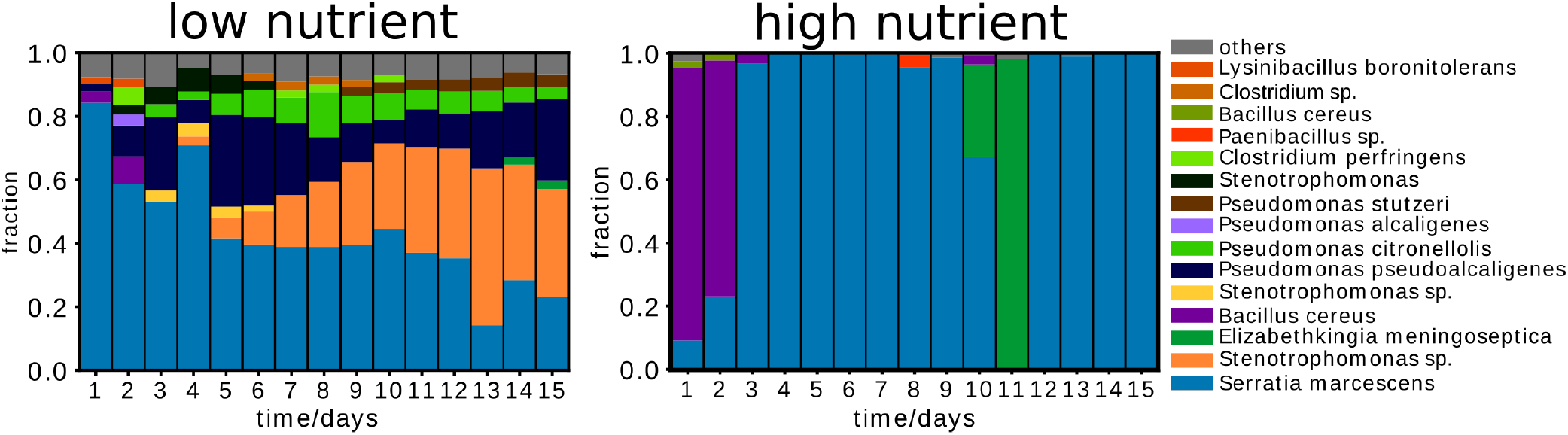
Nutrient concentrations impact dynamics and composition of a soil microbiota. Typical time-course of the community composition at low and high nutrient concentrations and thus weak and strong interactions according to Fig. 1. The plots show the change of composition over time based on 16S amplicon sequencing for a compost sample. Replicates from compost and other sampling sites (indoor flower pot and outdoor soil) show similar dynamics as shown in Supplementary Fig. 9. The amount of eukaryotes in those microcosms is very low (Supplementary Fig. 10). We can also see that several of the species found in the complex communities were also used for the pairwise interaction experiments shown in Fig. 1 and are therefore good representative of these complex soil communities. The composition of the start communities (day0) are shown in Supplementary Fig. 8.

These time-courses reveal striking differences between the low and high nutrient conditions; at low nutrient concentrations there were more species present and the temporal change of the system was rather ‘smooth’ (compost community shown in Fig. 2, others in Supplementary Fig. 9). On the contrary, at high nutrient concentrations the community exhibited sudden jumps between several low diversity states.

To gain intuition into whether the properties of the microbial interactions found in mono and co-culture (Fig. 1) may explain the behavior of complex communities (Fig. 2), we developed a mathematical model in which bacteria interact by changing the environment and are at the same time affected by these environmental changes. The model is a multi-species extension of a model we previously used to understand homogeneous populations and pairwise interaction outcomes^20^.

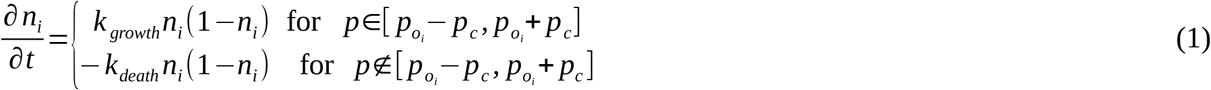

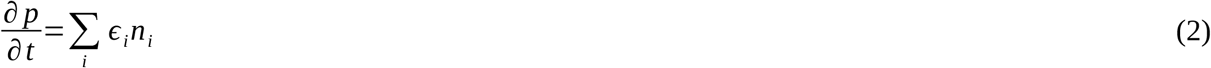

The bacterial species n_i_ grow logistically with growth rate k_growth_, but only if the environmental parameter p lies within the suitable range [p_oi_-p_c_, p_oi_+p_c_]. Outside that range the bacteria die with rate kdeath. Additionally, the bacteria change the environmental parameter p with rate *ϵ*_*i*_, which is taken from a uniform distribution in the interval [-c_p_, c_p_]. Accordingly, c_p_ is the maximal amplitude of the environmental change. At the end of every growth cycle the system is diluted with a constant factor (see Supplement for details).

Simulating 40 interacting pairs with this model and varying the extent to which they changed the environment and thus the interaction strength lead to results similar to what we observed experimentally (Fig. 3A purple, for more values of c_p_ see also Supplementary Fig. 17). Increasing the modification of the environment (c_p_) led to a loss of coexistence in co-culture, as seen in the experiments (Fig. 1D and Fig. 3B, violet). Since this model recapitulated the findings for pairwise interactions we were curious what it could tell us about complex communities. For this purpose, the above simulations were repeated with communities containing 20 species. Increasing the environmental modification by the bacteria caused a drop of biodiversity (Fig. 3A), which is in line with similar findings in Lotka-Volterra models^8^.

**Figure 3:**
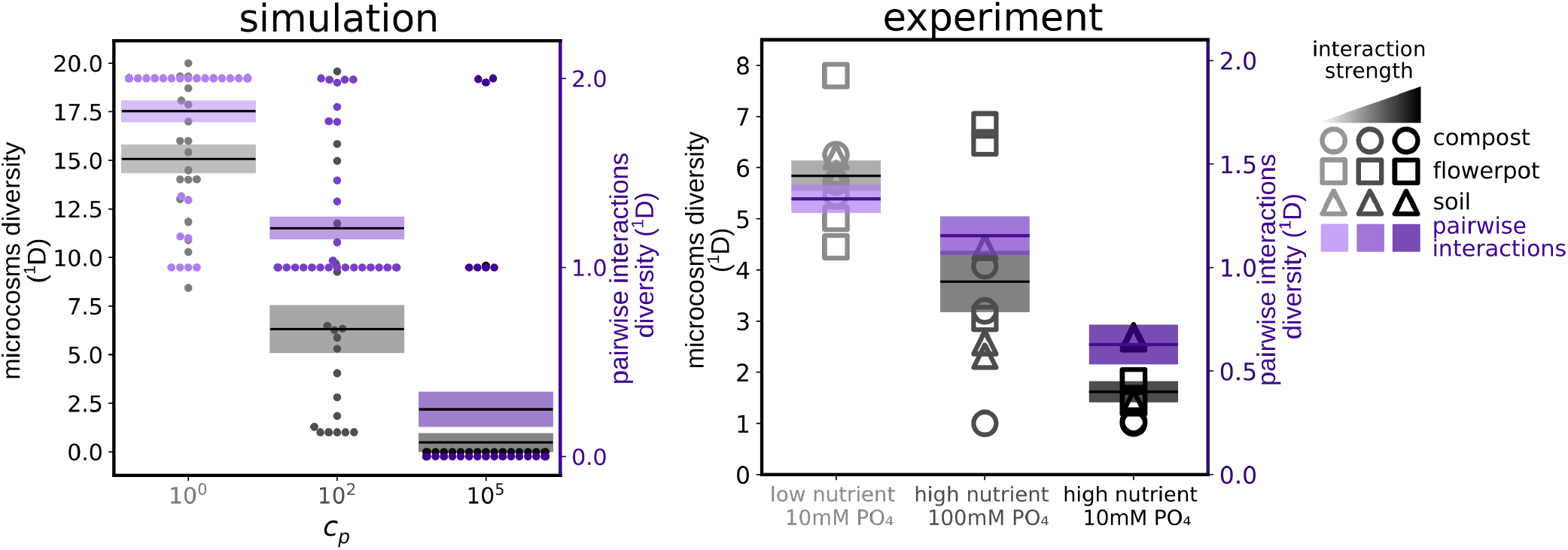
Increased interaction strength leads to a loss of biodiversity. (A) Simulations show a loss of coexistence in pairwise interactions (purple) and a loss of biodiversity in complex communities (20 interacting species, grey) upon increasing the strength by which the bacteria change the environment and thus interact. (B) The same behavior can be found in the experiments, where an increase in nutrient concentrations leads to a loss of diversity in both pairs as well as complex communities. Adding 100mM phosphate buffer in those experiments reduces the loss of biodiversity. The pairwise interaction outcomes shown in purple correspond to the data of Fig. 1D.

To confirm that this predicted drop of biodiversity could also be observed in the experiments propagating various complex communities, we calculated the diversity of the microbial communities at the end of the experiment for low and high nutrient conditions. Indeed, we observed a loss of biodiversity when the nutrient concentrations and thus the interaction strength was increased, as predicted by the model (Fig. 3B). pH modification could be identified as an important driver for the pairwise interactions in Fig. 1 (Supplementary Fig. 1,2 and 5). Accordingly, adding buffer to the complex communities also reduced the loss of biodiversity in high nutrient conditions. Therefore, the loss of biodiversity was largely driven by modifications of the environmental pH, not by the loss of limiting resources upon adding nutrients^9^. Overall, high nutrient concentrations caused stronger environmental modifications and interactions, leading to a loss of biodiversity in the microbial communities, as predicted by our simple model.

Another important property of ecosystems that seems to be linked to biodiversity is their stability, eg how unchanged an ecosystem remains over time^7,9,14^. We show and discuss in the following how interaction strength impacts the stability of the complex microcosms (the effects on pairwise interactions are similar and can be seen in Supplementary Fig. 11). To get an impression of how interaction strength might affect stability of microbial communities, we performed simulations with the above model to obtain the total bacterial density (∑n_i_) over time for weak and strong interactions, eg weak and strong modification of the environment (c_p_). Our model predicts that the fluctuations of the total bacterial density were much higher at stronger interactions (Fig. 4A, top).

**Figure 4:**
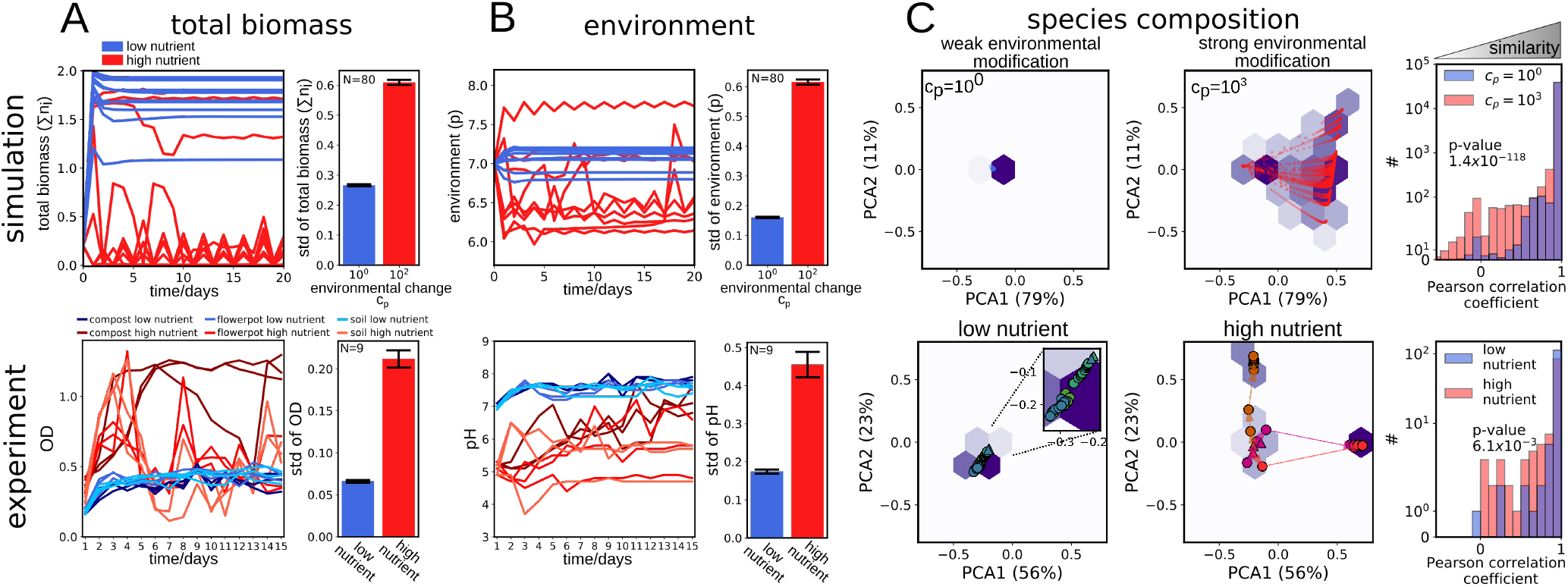
Stronger interactions lower stability of total biomass, environment and species composition. Data shown in red corresponds to high nutrient concentrations (strong interaction) and data in blue to low nutrient concentrations (low interaction). (A) Total bacterial density fluctuates more over time for stronger interaction in both the simulation (upper panel) and experiment (lower panel). On the left several example time curves are shown whereas the bar plots on the right show the mean of the standard deviations for all obtained time curves. (B) Also the environment fluctuates stronger for stronger interactions in the model (upper panel) and the experiments (lower panel). Again on the left example curves are shown and the mean of the standard deviations for all curves are on the right. (C) For weak interactions the compositions of the communities stay similar (upper left, simulation, lower left, measurement) over time whereas for strong interaction strength more pronounced changes in composition over time can be observed (upper middle, simulation; lower middle, measurement). The left and middle panel show example curves (different colors correspond to different replicates, arrows point into direction of time, triangles indicate day 1, data of the remaining samples is shown in Supplementary Fig. 15 and of simulations in Supplementary Fig. 22). The right panels show Pearson correlation coefficients of the composition between subsequent days for all obtained data. The closer the Pearson correlation coefficient to one the more similar are the compositions of two subsequent days, eg at stronger interactions the communities are more dissimilar between days. Simulation and measurement outcomes for multiple interaction strengths are shown in Supplementary Fig. 13, 14, 18 and 19.

To determine if this predicted loss of stability was present in our experimental communities, we analyzed the total biomass over time (as quantified by optical density). Consistent with our model predictions, we found that high nutrient concentrations caused stronger temporal fluctuations in all samples (Fig. 4A, bottom). In addition to increased fluctuations of the total bacterial density, the model predicted an increase in fluctuations of the environmental parameter p would show stronger fluctuations at stronger interactions (Fig. 4B, top). Consistent with this prediction, we found the same effect in the experiments when the pH, as a central environmental parameter, was measured over time (Fig. 4B, bottom). Finally, looking at the change of the bacterial composition, the model predicted stronger fluctuations of the composition over time at higher interaction strength, which again could be found in the measurements (Fig. 4C). We therefore found that stronger interactions led to a loss of the stability of total biomass, environment, and species composition as predicted by the model.

## Discussion

Despite its fundamental importance in ecology—and its current decline around the world^24^—a clear understanding of what determines biodiversity is still missing^6,25^. Abiotic factors surely influence biodiversity, but also interactions between organisms are thought to play a major role in setting the biodiversity of ecosystems^8,25,26^. However, how exactly interspecies interactions influence community diversity remains unclear since it is difficult to measure these interactions, and even more difficult to manipulate them experimentally. We showed here a way to tune the interaction strength between bacteria, which allowed us to understand how interactions set the biodiversity of microbial communities. High nutrient concentrations caused stronger microbial interactions, which led to less diverse communities.

This diversity loss is reminiscent of eutrophication, an over-enrichment of ecosystems with nutrients that often leads to a drastic loss of biodiversity^27^. Also, for eutrophication a stronger competition between species at increased nutrient concentrations – eg by limiting light – was suspected to contribute to biodiversity loss^28^. This raises the possibility that eutrophication processes could impact a wide range of different microbial communities. Such eutrophication may even be medically relevant. In the human gut microbiome, a loss of biodiversity was associated with western, high-caloric and low complexity diets compared to fiber rich, low caloric nutrition^29,30^. We speculate that such a loss of biodiversity upon easily accessible nutrients may be driven by an increased interaction strength between the gut microbes.

There exists a variety of evidence for the connection between biodiversity and stability. Higher biodiversity often – but not always - comes with higher stability in ecosystems^7,9,11,12,14,31^. In our experiments the increased interaction strength decreased stability in pairwise co-cultures as well as in complex communities, indicating that the loss of stability was independent of the actual biodiversity of the microbial system. The loss of stability seems therefore not to be directly caused by the biodiversity itself, but the interaction strength between the organisms negatively affects both biodiversity and stability at the same time.

Using simple microbial systems in the lab with the goal to investigate basic principles of ecology and evolution has lead to many fundamental insights^32,33^. However, because of the simplicity of those systems it is often rather unclear how far the obtained findings can be transferred to natural, much more complex communities. We show here that at least biodiversity and stability of complex systems can be understood from properties of simple pairwise interactions. For these ecosystem properties, the mean interaction strength seems to be more important than how the specific interaction pairs sum up to build the community. This surprising simplicity suggests that it is possible to not only understand complex microbial communities, but ultimately to engineer them.

## Methods

### Media, buffer and bacterial culture

All chemicals were purchased from SigmaAldrich (St.Lous, USA) unless stated otherwise.

Pre-cultures of bacteria were made in 1xNutrient medium (10g/l of yeast extract and 10g/l of soytone (both Becton Dickinson, Franklin Lakes, USA), 100mM Sodium phosphate, pH7), or Tryptic Soy Broth (Teknova, Hollister, USA) called TSB in the following. The experiments were performed in Base medium which contained 1g/l yeast extract, 1 g/l soytone, 0.1 mM CaCl_2_, 2 mM MgCl_2_, 4 mg/l NiSO_4_ and 50 mg/l of MnCl_2_. Different amounts of phosphate, glucose and urea were added depending on the experimental conditions as outlined below. The initial pH was adjusted to 7 unless stated otherwise. All media were filter sterilized using VWR Bottle Top Filtration Units (VWR, Radnor, USA). For plating of bacteria the cultures were diluted in phosphate buffered saline (PBS, Corning, New York, USA). Plating was done on Tryptic Soy Broth agar, with 2.5% agar (Becton Dickinson, Franklin Lakes, USA). For the experiments the bacteria were grown in 96-deepwell plates (Deepwell plate 96/500μL, Eppendorf, Hamburg, Germany) covered with AearaSeal adhesive sealing films (Excell Scientific, Victorville, USA). The growth temperature was 30°C for the isolates and 25°C for the complex communities, unless stated otherwise. The deepwell plates were shaken at 1350 rpm shaking speed on a Heidolph Titramax shakers (Heidolph, Schwabach, Germany). To avoid evaporation the plates were incubated inside custom build acrylic boxes. The exact conditions are outlined for the single experiments below.

### Estimation of population density (CFU/ml)

For CFU counting the bacteria were either added as droplets on the agar surface of 150mm petri dishes (droplet plating) or fully spread on 100mm agar plates (spread plating). The first method gives a high throughput since 96 cultures can be plated in one working step, but the second gives a higher accuracy in counting.

#### 1) Droplet plating

The cultures of interest were serially diluted in PBS (PBS; Corning, New York, USA) by seven 1/10-fold dilutions (20μL into 180μL, maximal dilution 10^−7^ x) with a 96-well pipettor (Viaflo 96, Integra Biosciences, Hudson, USA) using the program *“pipet/mix”* (pipetting volume: 20μl, mixing volume: 150μl, mixing cycles: 5, mixing and pipetting speed: 8). 10μl of every well were transferred on a large (150-mm diameter) Tryptic Soy Broth 2.5% agar plate (Tryptic Soy Broth (Teknova, Hollister, USA), Agar (Becton Dickinson, Franklin Lakes, USA)) with the 96-well pipettor (program *“reverse pipette”*, uptake volume: 20μl, released volume: 10μl, pipetting speed: 2). Droplets were allowed to dry in and the plates were incubated at 30°C for one to two days until colonies were visible. The different dilution steps allowed to find a dilution at which colonies could be optimally counted (between ~5 and ~50 colonies).

#### 2) Spread plating

The cultures were diluted in PBS with 7x 1/10x dilutions as described above and 150μL of the 10^−2,^ 10^−4^ and 10^−6^ dilutions were spread onto 100mm TSB agar plates with glass beads. The different dilutions again allowed to find a plate with optimal density for colony counting.

### pH measurement

To measure the pH of the microbial cultures, 170μl of sample were transferred into 96-well PCR plates (VWR, Radnor, USA) and the pH was measured with a pH microelectrode (Orion PerpHecT ROSS, Thermo Fisher Scientific, Waltham, USA).

### Measuring pH change of soil isolates

The soil isolates were isolated from local soil (Cambridge, MA, USA) as described elsewhere ^20,34^. The bacteria were pre-cultured in 1x Nutrient medium for 24h at 30°C. The cultures were diluted 1/100x into 200μL of

- Base, 10mM PO_4_, pH7
- Base, 10mM PO_4_, 1% glucose, 0.8% urea, pH 7
- Base, 10mM PO_4_, 0.4% glucose, 0.32% urea, pH 7
- Base, 100mM PO_4_, 1% glucose, 0.8% urea, pH 7

The bacteria were grown in these media for 24h at 30°C. Afterwards the pH was measured. The bacterial density was measured as optical density at 600nm (OD_600nm_) in 100μL in 96-well flat bottom plates (Falcon, Durham, USA) and only those pH values were taken into final consideration for which the corresponding culture reached on OD of at least 0.04. The results of the first two media conditions can be seen in Fig. 1 all results are shown in Supplementary Fig. 2.

### Measuring bacterial growth in spent media

8 soil species were chosen for this experiment: Pseudomonas putida (ATCC#12633), Pseudomonas aurantiaca (ATCC#33663), Pseudomonas citronellolis (ATCC#13674), Micrococcus luteus (Ward’s Science, Rochester, NY), Sporosarcina ureae (Ward’s Science, Rochester, NY), Bacillus subtilis (strain 168), Enterobacter aerogenes (ATCC#13048), Serratia marcescens (ATCC#13880). Those species can be differentiated by colony morphology (Supplementary Fig. 3) and have been used for interaction studies before^18,35^. The bacteria were grown in 5mL TSB (Teknova, Hollister, USA) overnight at 30°C. The bacteria were spun down (15mins, 3220g, Eppendorf Centrifuge 5810) and re-suspended in 5mL Base medium. The washed bacteria were diluted 1/100x into 2x 5mL Base, +/− 1% glucose, 0.8% urea, with either 10mM or 100 mM phosphate, pH7 (spent media cultures). At the same time a new pre-culture was set up in TSB as described above. Both cultures were grown for 24h at 30°C. The spent media cultures were spun down (15mins, 3220g, Eppendorf Centrifuge 5810) and the supernatant filter sterilized with a 50mL Steriflip Filtration Unit (SCGP00525, 0.22μm, Millipore/SigmaAldrich, St. Louis, USA). 50μL of this spent media were spotted onto Tryptic Soy Agar plates to verify sterility. The spent media were either used directly or supplemented with 1/20x of 20x original media without phosphate buffer to replenish the nutrients. The second pre-culture was spun down as well after 24h (15mins, 3220g, Eppendorf Centrifuge 5810) and re-suspended with base medium as described above. Those bacteria were now diluted 1/100x into the spent media and also into the corresponding fresh media that are described above. The cultures were grown 24h at 30°C in 96-deepwell plates (Deepwell Plate 96/500 μl, Eppendorf, Hamburg, Germany) 200μL per well in shaken culture (1350 rpm shaking speed on a Heidolph Titramax shaker). After 24h the OD_600nm_ of the cultures (100μL in 96-well flat bottom plates (Falcon, Durham, USA)) in the different spent media was measured and divided by the OD_600nm_ obtained in fresh media. The resulting data is shown in Fig. 1C and Supplementary Fig. 4 and 5.

### Pairwise interactions

The 8 soil strains described above were grown in TSB overnight at 30°C. The bacteria were spun down 5mins at 3220g in an Eppendorf centrifuge 5810 and resuspended in 2.5mL base medium, with 10mM Phosphate, pH7. For each of the 28 pairwise combinations 10μL of each strain were diluted into 200μL Base, 10mM/100mM PO_4_, +/− (1% glucose, 0.8% urea). The co-cultures were incubated at 30°C and 1350rpm shaking speed on a Heidolph Titramax shaker in 96-deepwell plates. Every 24h the co-cultures were diluted 1/10x into fresh media. The pH and OD_600nm_ were measured at the end of every incubation cycle (every 24h). After 5days the co-cultures were plated by droplet plating as described above. The agar plates were incubated at 30°C for around 2 days until colonies were clearly visible. The colonies were then counted. The ^1^D diversity was calculated according to 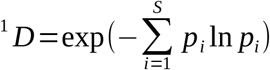, where ^1^D was set to 0 if both species went extinct. The results are shown in Fig. 1D and Supplementary Fig. 5.

### Obtaining environmental samples

The compost used for the experiments was purchased from Bootstrap Compost in Boston, Massachusetts. The soil was sampled in Cambridge, Massachusetts, at a depth of ~30 cm. The soil was kept at 4°C until the experiments were performed. Flower pot soil was sampled the day of the experiment by taking soil from a large plant pot at depth 10cm.

### Temporal dynamics of soil microcosms

For the compost and the flower pot experiments, 4g of sample were diluted in 20ml of PBS, vortexed at intermediate speed for 30s and incubated on a platform shaker (Innova 2000, Eppendorf, Hamburg, Germany) at 250rpm and room temperature. After 30 minutes, the samples were allowed to settle for 5 minutes and the supernatant was transferred to a new clean tube. The sample was then diluted 1:10 before inoculation of the experiments. For the soil experiment, 4 grains of soils (~0.1g) were diluted in 40ml of PBS, vortexed and mixed as described for the compost samples. The supernatant collected after settling was directly used for inoculation without further dilution. Experiments were inoculated by mixing 170μl of these obtained liquids into 1530μl of appropriate media as indicated below.

Experiments were performed in 2000-μl 96-deepwell plates (Deepwell Plate 96/2000 μl, Eppendorf, Hamburg, Germany) using Base media at pH 7 to which either 10mM (referred to as “low buffer”) or 100mM (referred to as “high buffer) phosphate were added. 0/0%, 0.5/0.4%, 1/0.8%, 2/1.6%, 3/2.4% and 5/4% of glucose/urea (m/V) were added to the high and low buffer media respectively. Plates were covered with two sterile AearaSeal adhesive sealing films (Excell Scientific, Victorville, USA) and incubated at 25°C on a VWR Micro Plate Shaker at 500 rpm.

Every 24 hours, the cultures were thoroughly mixed by pipetting up and down 30 times using the Viaflo 96-well pipettor (mixing volume: 300μl, speed: 10, cycles 30). Then the cultures were diluted 1:10 into fresh media. At the end of every cultivation day 170μl of culture were transferred into flat bottom 96-well plates (Falcon, Durham, USA) and the optical density (OD_600nm_) was measured with a Varioskan Flash (Thermo Fisher) plate reader. The pH was measured as described above. The remaining culture liquid was stored at −80°C for subsequent DNA extraction. The DNA extractions were performed using Agencourt DNAdvance A48705 extraction kit (Beckman Coulter, Indianapolis, IN, USA) following the provided protocol. The obtained DNA was used for 16S amplicon sequencing of the V4-V5 region. Some amount of the samples was also checked for eukaryotes by sequencing the 18S V4 region. The sequencing was done on a Illumina MySeq by CGEB - Integrated Microbiome Resource at the Dalhousie University, Halifax, NS, Canada.

### Data analysis

We analyzed the obtained 16S reads as described elsewhere^36^. From the 16S reads the amplicon sequence variants (ASVs) were obtained with dada2 package in R^37^. Taxonomic identities were assigned to the ASVs by using the GreenGenes Database Consortium (Version 13.8) ^38^ as reference database. The principle component analysis for Fig. 4 was performed with scikit-learn package in Python^39^.

## Author contributions

C.R., J.B. and J.G. designed the research. J.B. and C.R. carried out the experiments and performed the mathematical analysis. C.R., J.D. and J.G discussed and interpreted the results, and wrote the manuscript.

