## Supplementary Information and Data for "Strength of species interactions determines biodiversity and stability in microbial communities"

[2] Current address: Department of Molecular and Cellular Biology, Harvard University, Cambridge, MA, USA

* equal contribution

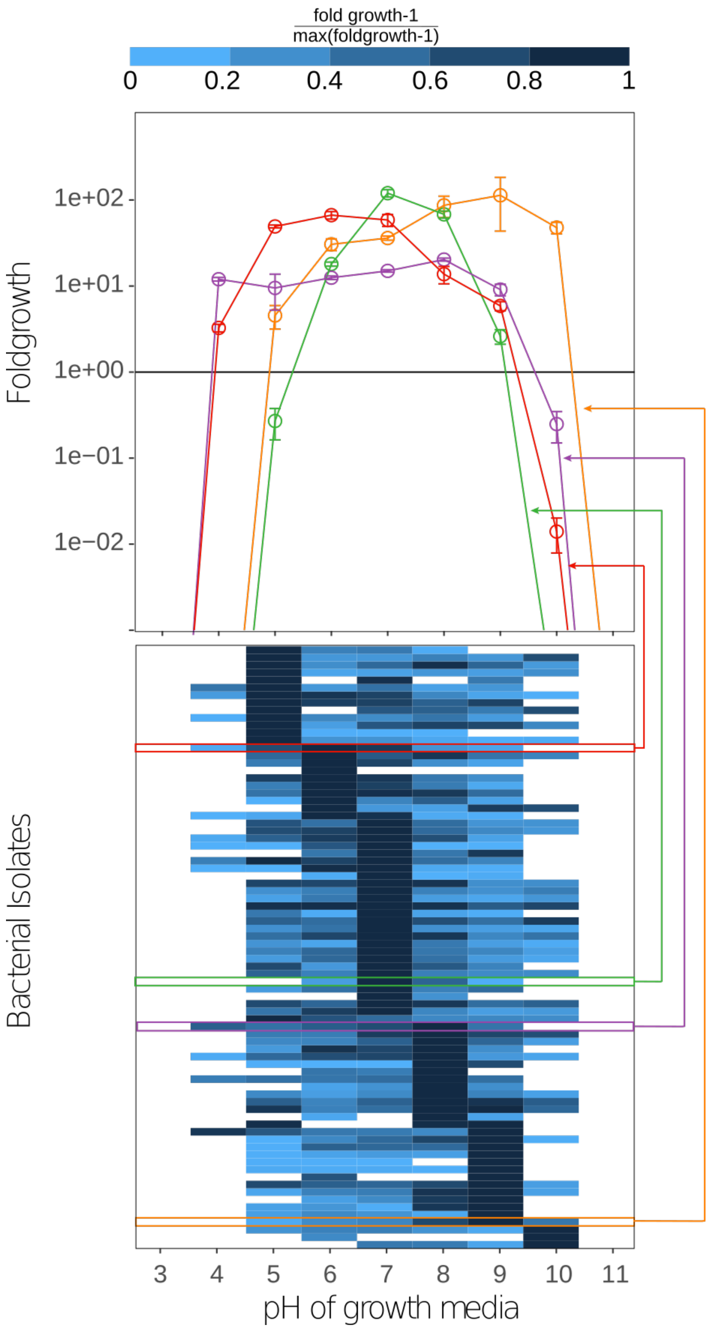


***Supplementary Figure 1: Different soil strains have different suitable pH ranges.*** *We tested the optimal growth pH of 81 isolated soil species. It is a subset of the species shown in Supplementary Fig. 2B. All isolates were pre-cultured in 200µL of 1xNutrient medium for 24h at 25°C with 1350 rpm shaking speed in 500-µl 96-deepwell plates (Eppendorf, Hauppauge, USA). After 24h of growth the cultures were diluted 1:100 into 500-µl 96-deepwell plates and a final volume of 200µl of Base media with 100mM phosphate with pH values of 3-11. Cultures were incubated for 24h at 25°C at 1350 rpm on a Heidolph Titramax shaker. Population densities were estimated by CFU counting at the start of the experiment and after 24h, which allows to estimate the fold growth in 24h that is shown in the figure. Several example curves are shown in the upper panel. As can be seen those curves can have several shapes. For simplification we decided to describe the shape of those curves with a heaviside function in our simulations (see below).*

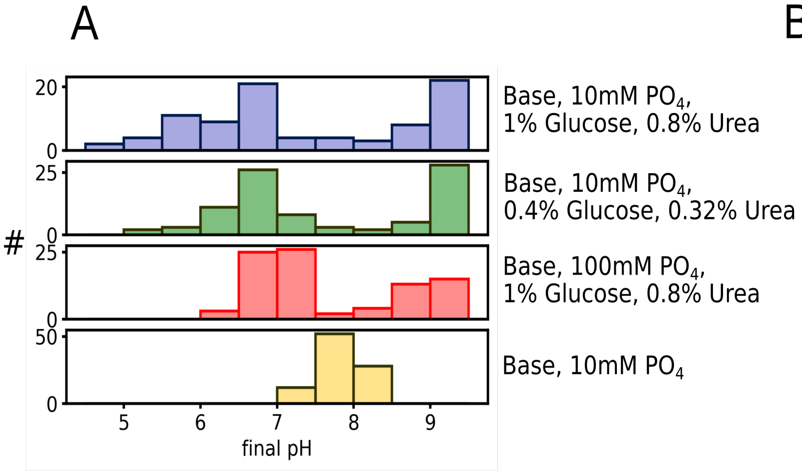


B

| Genus | low nutrient | high nutrient | Genus | low nutrient | high nutrient | Genus | low nutrient | high nutrient |
|---|---|---|---|---|---|---|---|---|
| Bacillus | 7.5 | 6.8 | Caryophanon | 7.7 | 7.8 | Paenibacillus | 7.4 | 4.9 |
| Rummeliibacillus | 7.1 | 7.2 | Bacillus | 8 | 6.5 | Bacillus | 7.8 | 5.8 |
| Bacillus | 7.8 | 9 | Lysinibacillus | 8 | 9.2 | Bacillus | 7.8 | 5.7 |
| Bacillus | 7.9 | 6 | Bacillus | 8.1 | 6.2 | Bacillus | 7.8 | 9 |
| Bacillus | 8 | 5.5 | Bacillus | 7.8 | 6.7 | Bacillus | 8 | 5.5 |
| Bacillus | 7.9 | 6.8 | Bacillus | 7.3 | 5.1 | Arthrobacter | 8 | 9.2 |
| Bacillus | 7.7 | 5.9 | | 8.1 | 9.1 | Arthrobacter | 8 | 6.8 |
| Bacillus | 7.8 | 9.1 | Bacillus | 8 | 5.5 | Arthrobacter | 8 | 9 |
| Bacillus | 7.8 | 5.5 | Paenibacillus | 7.3 | 4.6 | Arthrobacter | 8 | 7.3 |
| Bacillus | 7.6 | 9.1 | Paenibacillus | 7.2 | 6 | Bacillus | 7.9 | 6.9 |
| Zhihengliuella | 7.4 | 9.1 | Bacillus | 8 | 6.7 | Bacillus | 8 | 5.8 |
| Bacillus | 7.7 | 6.7 | Lysinibacillus | 7.7 | 7.8 | Bacillus | 7.9 | 6.7 |
| Bacillus | 7.8 | 6.9 | Bacillus | 7.7 | 9.1 | Bacillus | 7.9 | 6.6 |
| Bacillus | 7.9 | 7 | Bacillus | 7.9 | 6.6 | Paenibacillus | 7 | 5.8 |
| Arthrobacter | 8 | 8.5 | | 8 | 9 | Paenibacillus | 7.3 | 8.3 |
| Bacillus | 7.8 | 6.7 | Bacillus | 7.9 | 6.8 | Bacillus | 7.6 | 6.9 |
| Bacillus | 8 | 5.1 | Bacillus | 7.9 | 8.9 | Bacillus | 7.8 | 9.1 |
| Bacillus | 7.9 | 8.6 | Bacillus | 7.8 | 7 | Bacillus | 7.9 | 6.9 |
| Arthrobacter | 7.9 | 6.3 | Arthrobacter | 8 | 9.1 | Bacillus | 7.9 | 6.1 |
| Bacillus | 7.9 | 7 | Arthrobacter | 8.1 | 6.9 | | 7.9 | 8.7 |
| Bacillus | 7.9 | 5.4 | Bacillus | 7.6 | 6.5 | Arthrobacter | 8 | 9.1 |
| Bacillus | 7.8 | 8.6 | Bacillus | 7.9 | 6.2 | Bacillus | 8 | 5.6 |
| Bacillus | 7.8 | 6.6 | | 7.9 | 8.3 | | 8 | 9.1 |
| Bacillus | 7.8 | 9.1 | Bacillus | 7.8 | 8.1 | | 7.9 | 8.9 |
| Bacillus | 8 | 6.3 | | 7.9 | 7.9 | Planococcacea | 7.3 | 8.7 |
| Bacillus | 8 | 5.8 | Bacillus | 8 | 8.8 | Bacillus | 7.4 | 6.5 |
| Bacillus | 7.9 | 5.4 | Bacillus | 7.8 | 9.1 | | | |
| Bacillus | 7.9 | 6.5 | Bacillus | 7.9 | 9.2 | | | |
| Arthrobacter | 8 | 8.9 | Bacillus | 8 | 5.5 | | | |
| Paenibacillus | 7.2 | 4.6 | Bacillus | 8 | 6.3 | | | |
| Bacillus | 8 | 6.2 | Bacillus | 7.8 | 9.2 | | | |
| Bacillus | 7.8 | 9.1 | Bacillus | 7.9 | 9.1 | | | |

***Supplementary Figure 2: Nutrient concentrations and buffering determine pH change of growth media.*** *(A) The top and bottom panels show the same data as Fig. 1B. Using intermediate nutrient concentrations also causes intermediate pH shifts (green) compared to high (blue) and low (yellow) nutrient concentrations. Also adding higher concentrations of buffer lowers pH shifts (red) compared to the situation with low buffer (blue). (B) List of soil isolates that were used to measure the data in main text Fig. 1B and Supplementary Fig. 1A and 2B. Strains were identified down to genus level by sequencing their 16S rRNA gene and comparing it to the RDP database. The strains belong to a collection of soil strains that we used before for interaction studies [1,4]. As can be seen many of those strains belong the genus Bacillus, nevertheless they can change the pH into alkaline or acidic directions. For some cases the sequencing failed which lead to empty entries.*

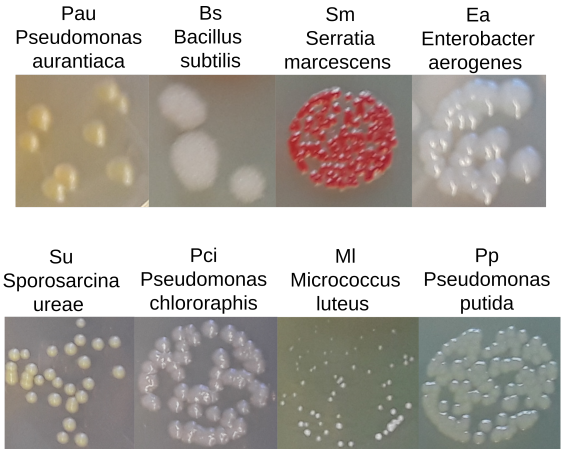


***Supplementary Figure 3: Bacteria for the pairwise interaction experiments.*** *The different colony morphologies allowed to distinguish them after plating on agar plates.*

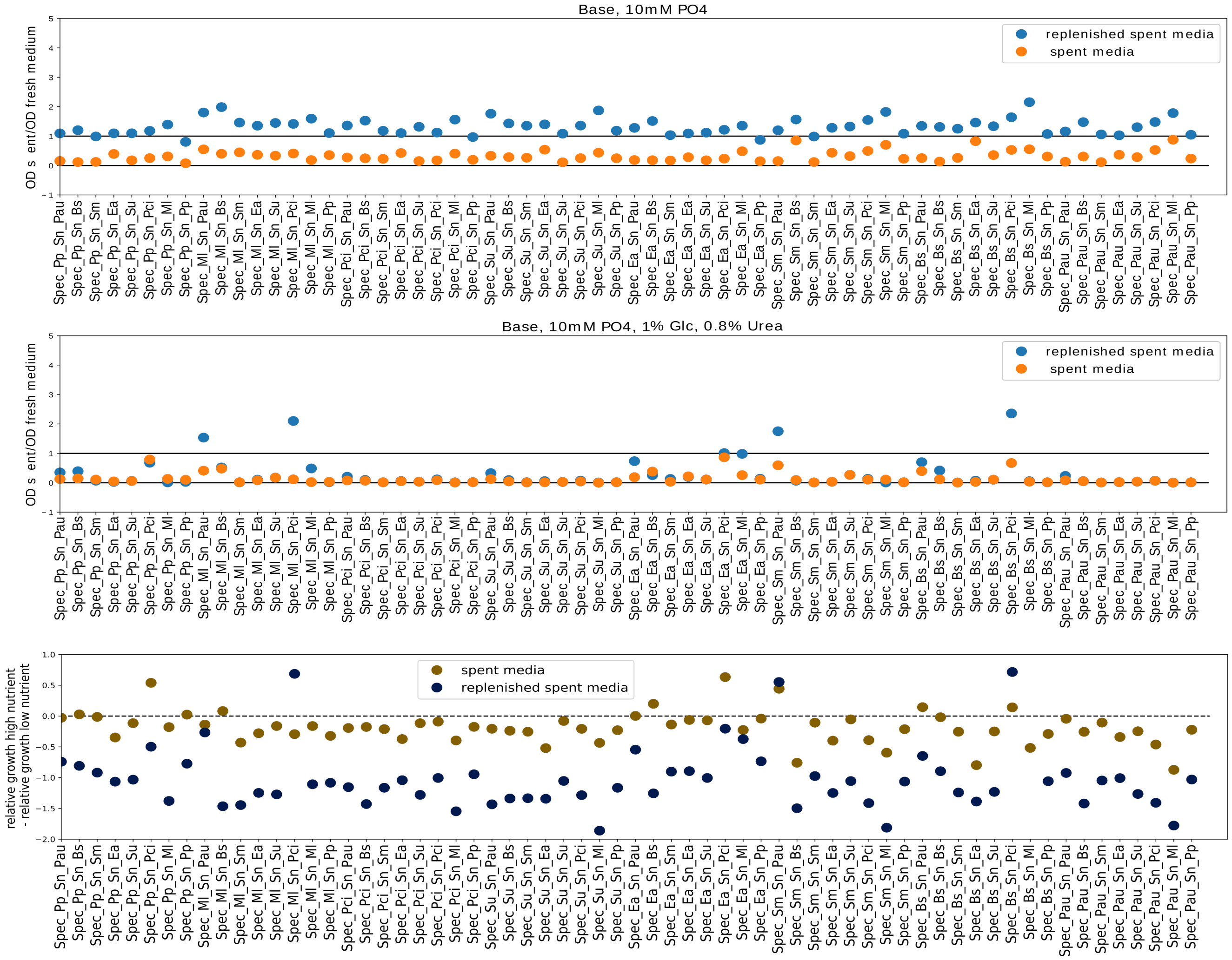


***Supplementary Figure 4: High nutrient concentrations lead to stronger negative interactions between bacteria.*** *The figure shows all the data of main text Figure 1C for low nutrient concentrations (top) and high nutrient concentrations (middle). The bottom panel shows the difference between the top and middle one. As can be seen in most cases (84%, for spent media without replenishment) increasing nutrient concentrations lead to a stronger inhibition of the interaction partner (values below zero), however in the remaining cases it leads to a relative facilitation (values above zero). Spec_X_Sn_Y means species X was grown in supernatant of species Y.*

*1.*

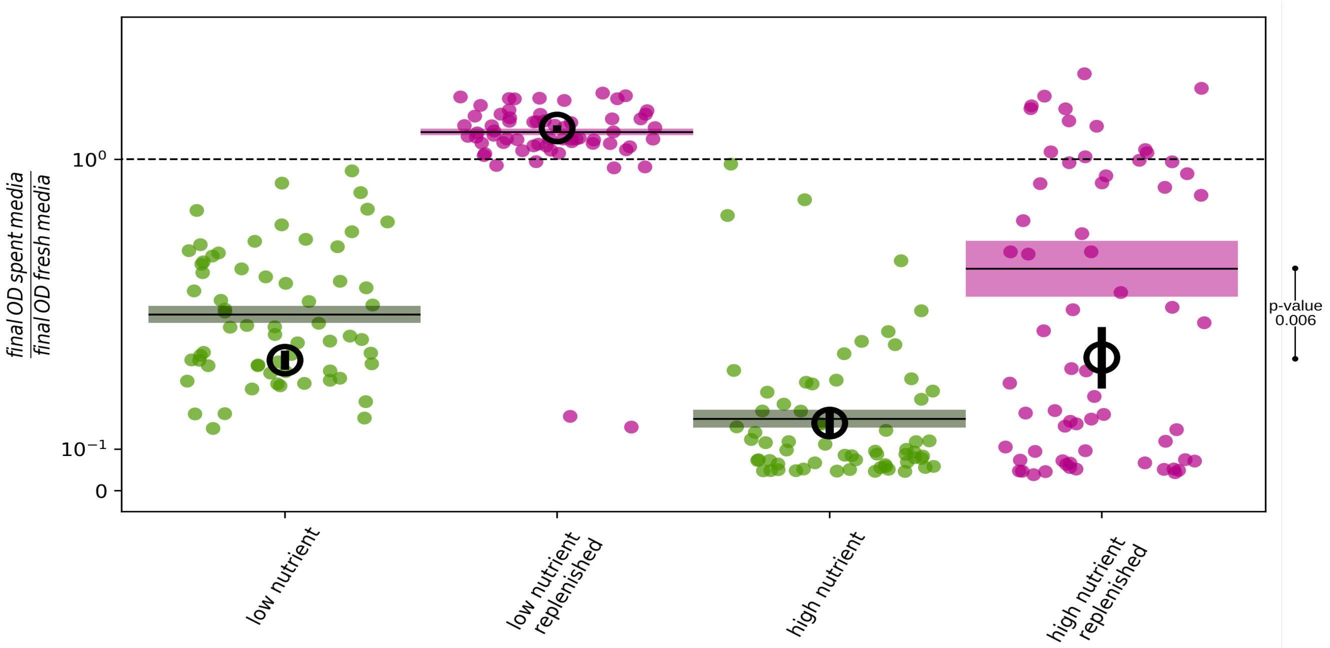


***Supplementary Figure 5: Growth inhibition caused by high nutrient spent media is partially caused by pH and can be removed by buffering.*** *The scatter plots show the ratio of final OD in spent and final OD in fresh media for all 64 interaction pairs in buffered media at low (left) and high (right) nutrient concentrations. The solid lines and boxes show the corresponding mean and SEM. This figure is thus equivalent to Fig. 1C in the main text with higher buffer concentrations (100mM $PO_4$). The black circles show the data of Fig. 1C eg with lower buffer concentrations (10mM $PO_4$). As can be seen the presence of higher buffer concentrations slightly facilitates growth in spent, but not replenished media, possibly because adding phosphate avoids phosphor to be a limiting resource. However, the strongest effect of buffering can be seen in the replenished supernatant. Whereas there is no effect upon the low nutrient replenished supernatant, bacteria grow much better in high nutrient replenished media with higher buffer concentration compared to lower phosphate (one-sided t-test p-value = 0.006). Since in the replenished media nutrient competition as a mode of interaction does not matter, this shows that the growth hindering and thus toxic effect of replenished high nutrient media can partially be diminished by buffering. Thus at least a part of the toxic effect of high nutrient supernatant is caused by pH.*

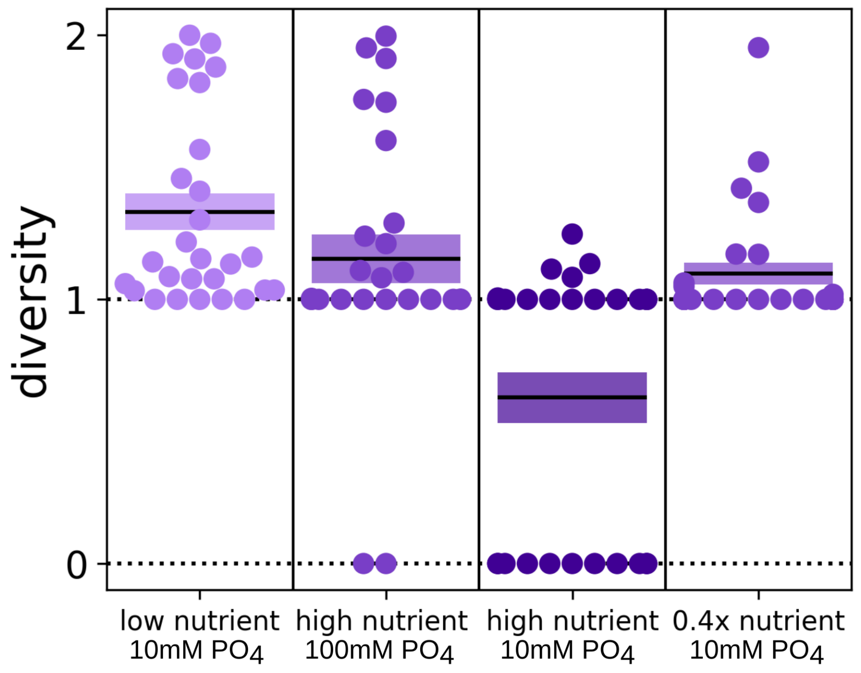


***Supplementary Figure 6: Nutrient levels determine interaction strength.*** *The first three columns correspond to Fig. 1D. The fourth column shows the interaction outcomes for a medium nutrient concentration of 0.4% glucose and 0.32% urea eg. 0.4x the high nutrient condition. As expected the results fall in between the results for the low (no Glucose and Urea) and high (1% Glucose and 0.8% Urea) nutrient outcomes.*

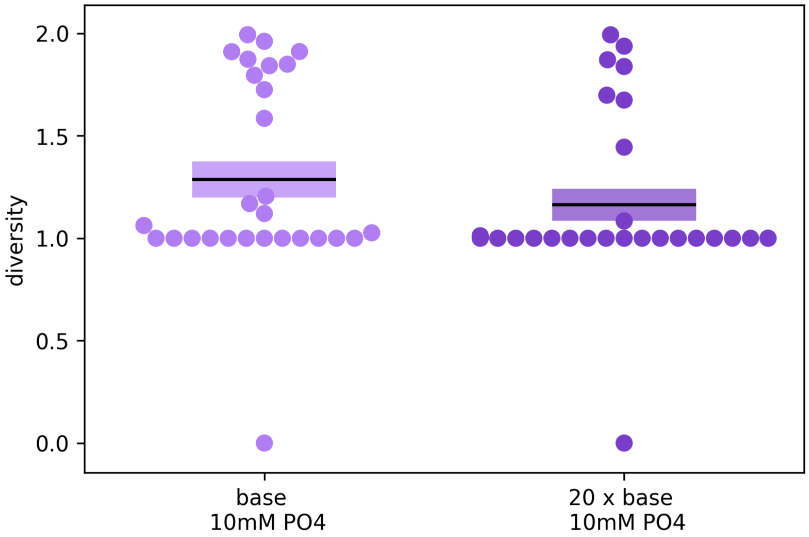


***Supplementary Figure 7: Complex nutrients weakly effect interaction.*** *Increasing the amount of yeast extract and soytone from 1g/L each to 20g/L leads to a slight decrease in overall diversity (p-value: 0.112). However, the effect of glucose and urea is much stronger. On reason for that may be that yeast extract and soytone also work as buffers, which stabilize pH at high nutrient concentrations.*

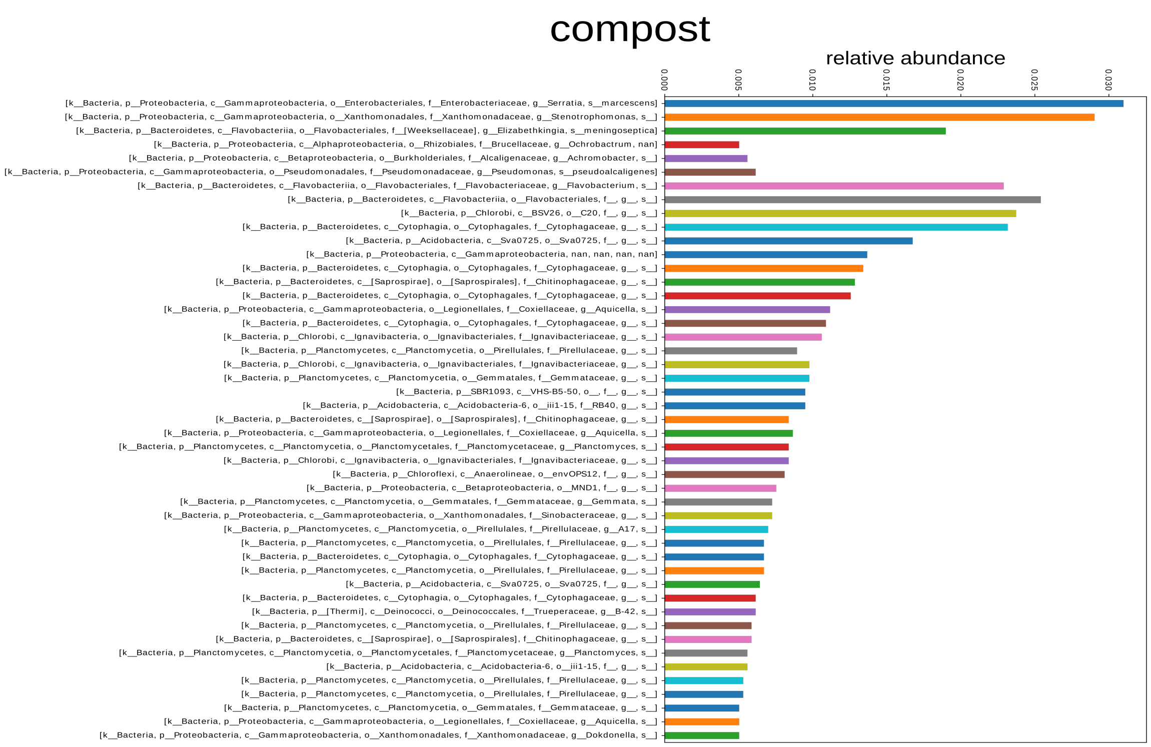


[1]D Diversity 193.8

Richness 330

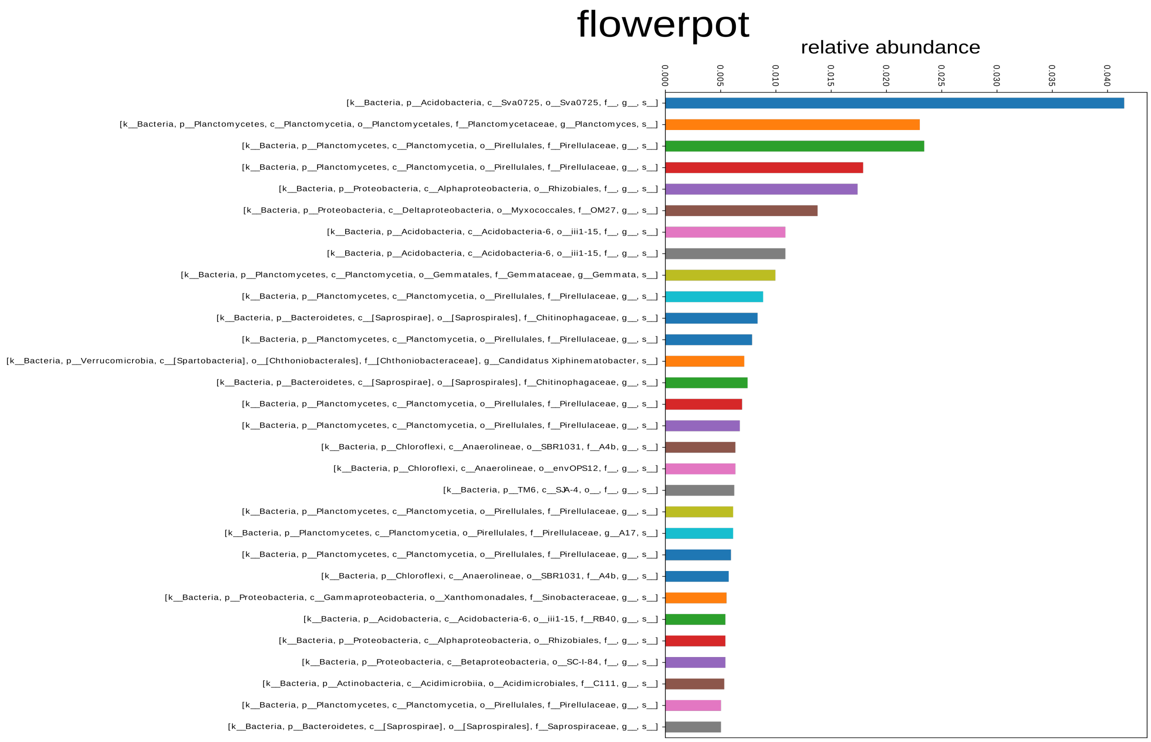


[1]D Diversity 347.8

Richness 625

***Supplementary Figure 8: Initial community compositions.*** *Shown are the ASVs with more then 0.05 abundance. The corresponding [1]D diversity and richness are much higher then at the end of the experiments (Fig. 3), eg those communities collapsed to communities with lower diversity during the experiments. The sequencing of the initial soil community failed.*

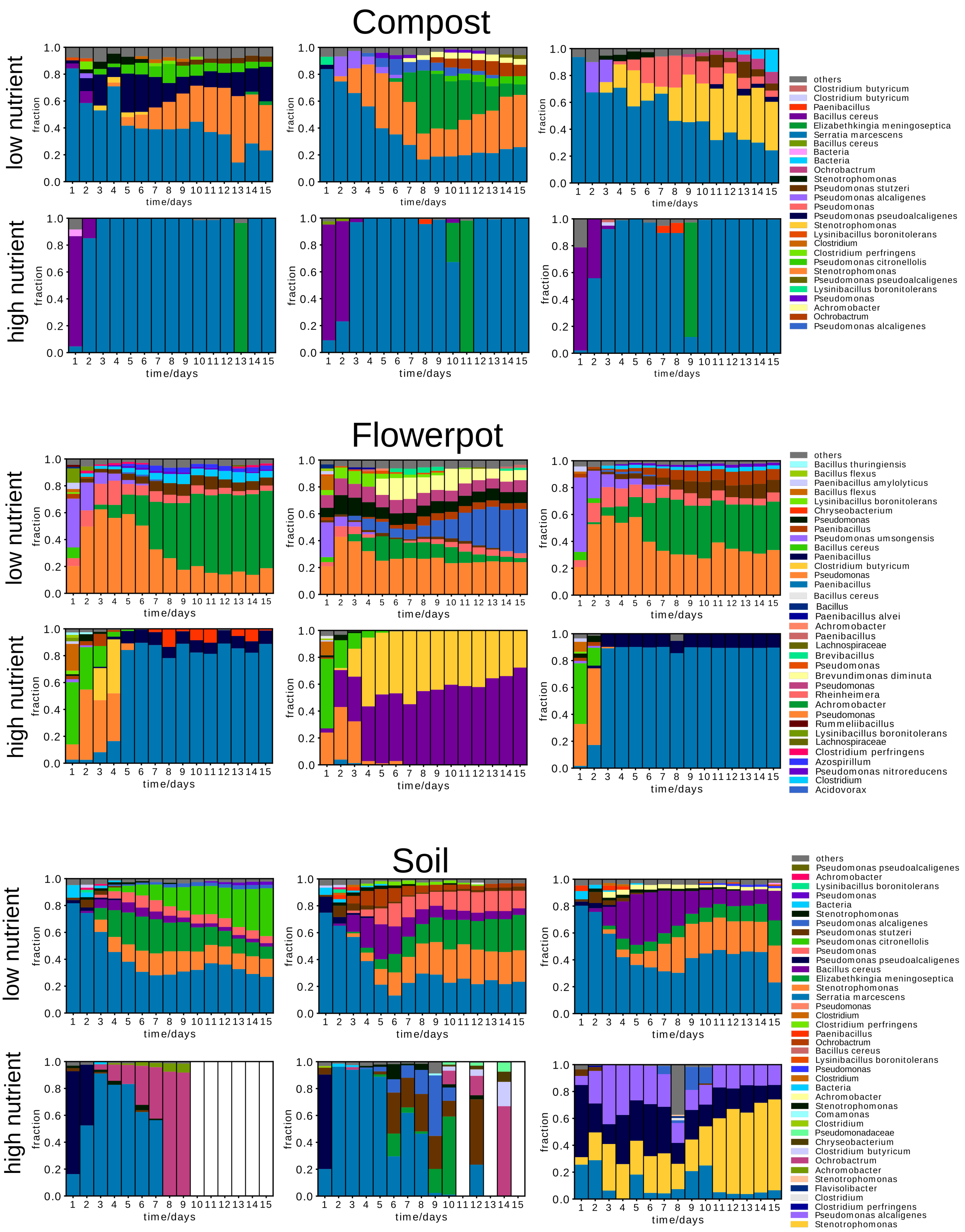


***Supplementary Figure 9: Community composition over time for different samples sites, replicates and nutrient conditions.*** *The colors that represent the different species are consistent for a specific sample (compost, flowerpot, soil), but may vary between them. In  a few cases different ASVs were identified as the same species, which causes a connection of the same species name with different colors within the same sample site. The white columns indicate days for which the sequencing failed.*

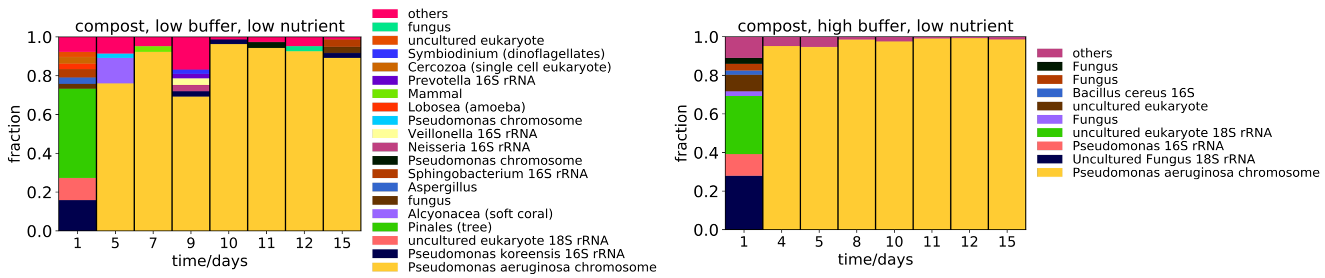


***Supplementary Figure 10: Eukaryotes in compost sample.*** *To see whether eukaryotes play a central role in these systems we also performed 18S rRNA sequencing on some of the samples. Many of them (43 out of 64) failed because of lack of DNA. Two time-courses that lead to some results are shown here, whereas several days are missing. As can be seen many of the found species are indeed not eukaryotes, but the primer amplified some bacterial DNA. Thus the overall amount of eukaryotes is rather low especially after the first day. We cannot exclude the presence of phages, but since the experimental outcomes can be changed by buffering the media they do not play a major role in the changing the dynamics of the systems.*

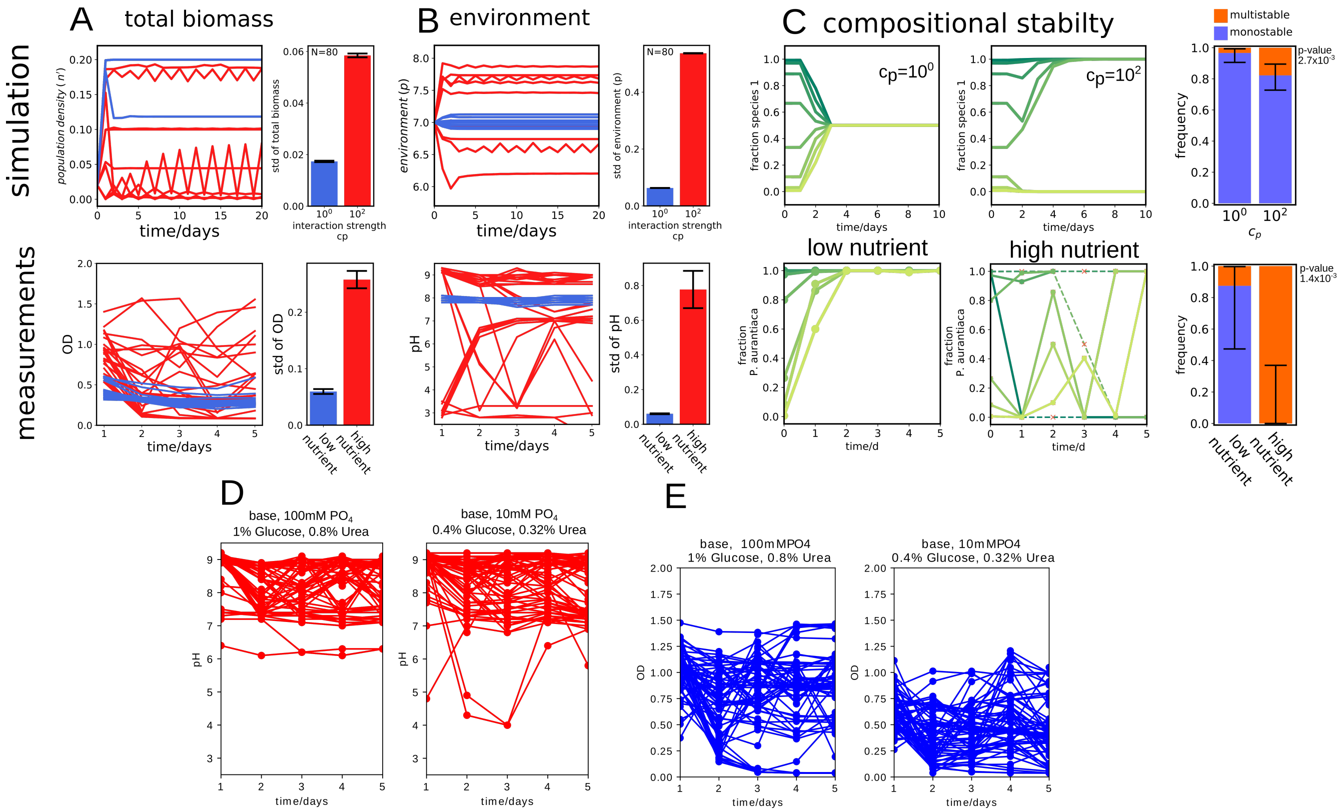


***Supplementary Figure 11: High nutrient concentrations lower stability of total biomass, environment and composition for pairwise interactions.*** *This figure is analogues to main text Fig. 4 but shows the effect of interaction strength on pairwise interactions instead of complex communities. (A) The total biomass fluctuates more over time upon high interaction strength (red) compared to weak (blue), both in the simulation (upper panels) as well as the experiment (lower panels). (B) Also the environment fluctuates stronger at higher interaction strength (red) then low interaction (blue). (C) The composition of the pairwise co-culture usually converges towards one fixed point for weak interactions, but shows more often bistability at high interaction strength in the simulations. The same situation can be found in the experiments, where at high nutrient concentrations the systems end up in different final states depending on the initial states in all tested cases. The amount of bistabilty is higher in the experiments than the simulation since in the experiments ecological suicide occurs more often then in the simulation. Ecological suicide itself is density depended and thus causes often multistability. The red crosses and dashed lines indicate interpolated values, when the CFU dropped below detection limit. Curves for all 8 tested interaction pairs can be found in Supplementary Fig. 12 and more simulated curves in Supplementary Fig. 20 and 21. Buffering the media or using lower concentrations of nutrients leads to smaller fluctuations of the pH (D) and less of the OD (E). p-values were calculated with Fisher's exact test. Errorbars in (C) are 95% Clopper-Pearson confidence intervals and SEM otherwise.*

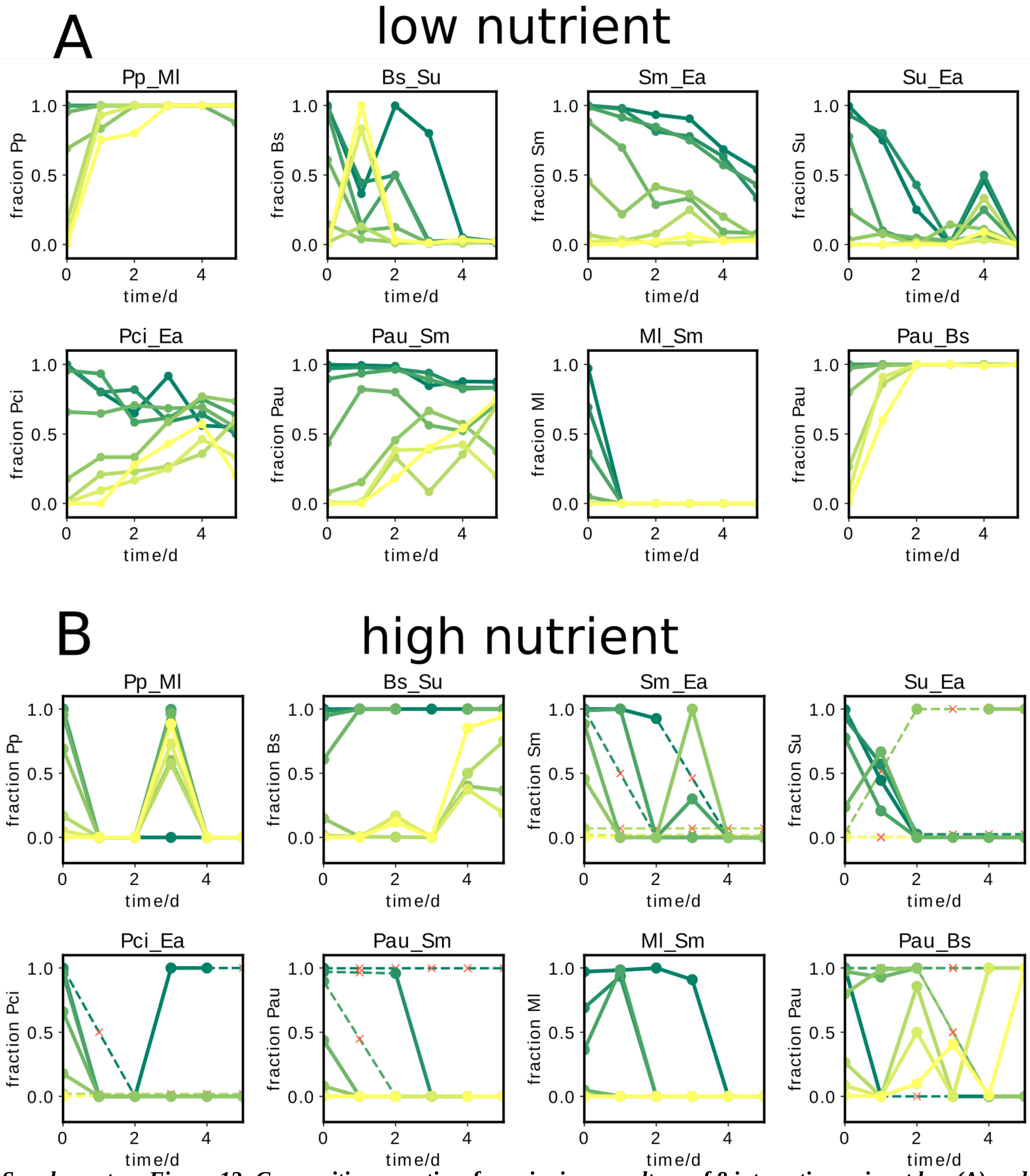


***Supplementary Figure 12: Composition over time for pairwise co-culture of 8 interaction pairs at low (A) and high (B) nutrient concentrations.*** *At low nutrient concentrations starting at different initial fractions of the interactions partners they usually converge towards similar final values after 5 days. (B) For high nutrient concentrations, the co-cultures show strong dynamics and end up with different values depending on the initial fractions. We call a system monostable when all initial fractions end up with species A or B dominating, both species coexists, or both species go extinct. Different outcomes for different fractions we call multi-stable. The red crosses and dashed lines indicate interpolated values, when the CFU dropped below detection limit*

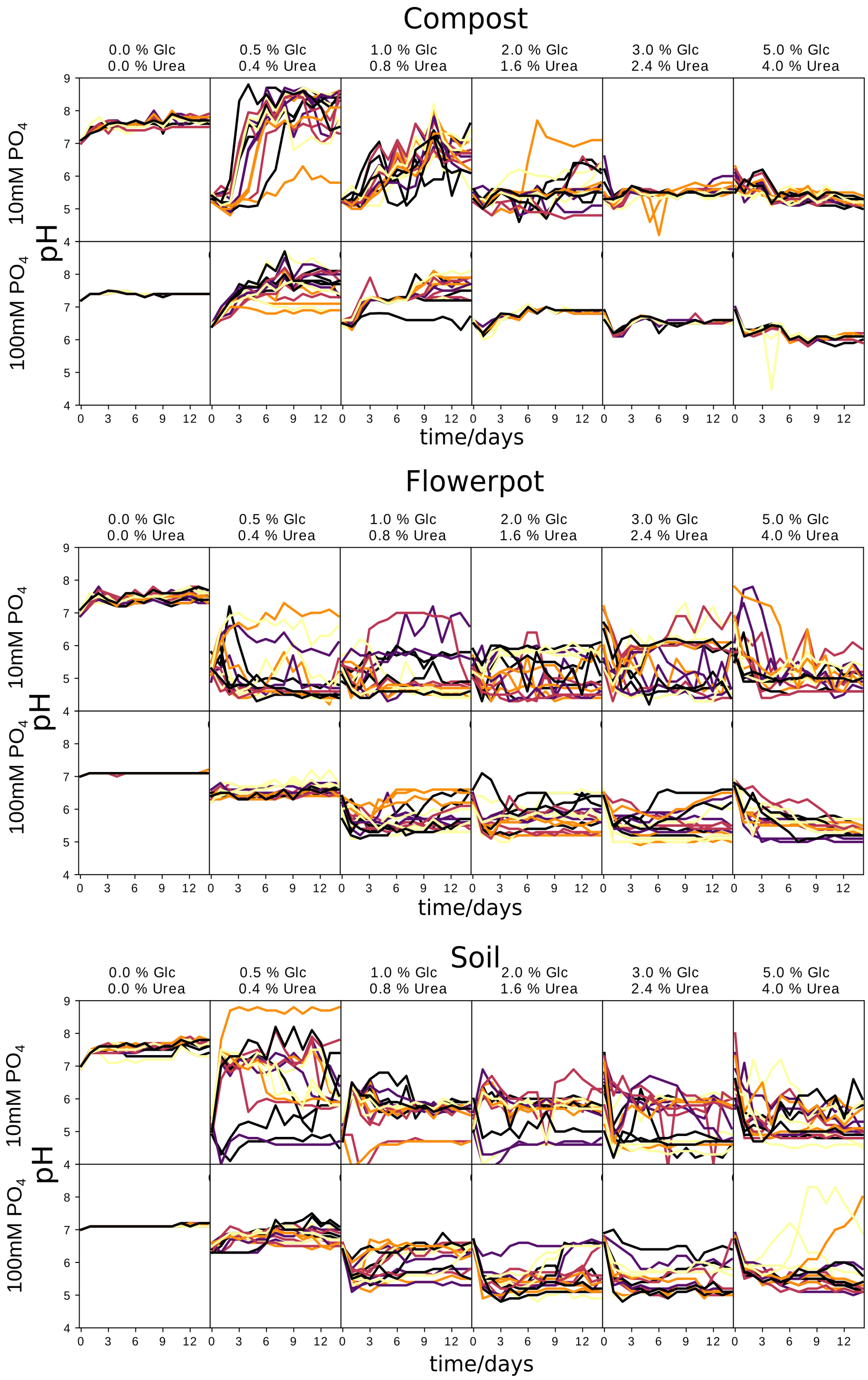


***Supplementary Figure 13: Fluctuation of pH depend on nutrient concentrations.*** *For each sampling site the upper row shows the situation for 10mM $PO_4$ buffer and the lower row for 100mM $PO_4$ buffer. The fluctuations increase first with the nutrient concentrations, but become for soil and compost smaller again at high nutrient concentrations, where likely the nutrient concentrations become so high that they become toxic themselves. The different curves show different replicates, there are 16 replicates for every condition.*

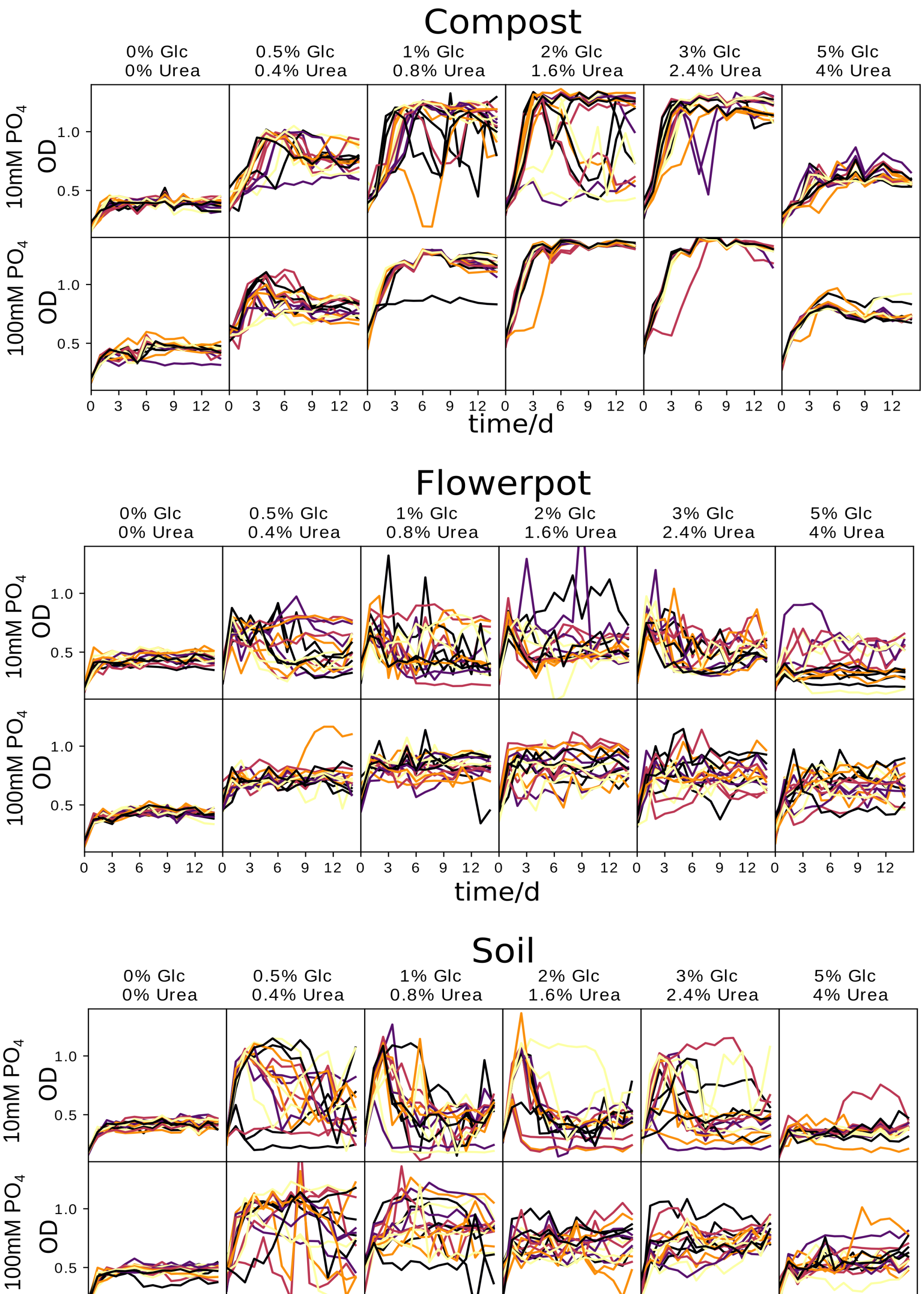


***Supplementary Figure 14: Fluctuation of OD depend on nutrient concentrations.*** *For each sampling site the upper row shows the situation for 10mM $PO_4$ buffer and the lower row for 100mM $PO_4$ buffer. The fluctuations increase first with the nutrient concentrations, but become for soil and compost smaller again at high nutrient concentrations, where likely the nutrient concentrations become so high that they become toxic themselves. The different curves show different replicates, there are 16 replicates for every condition.*

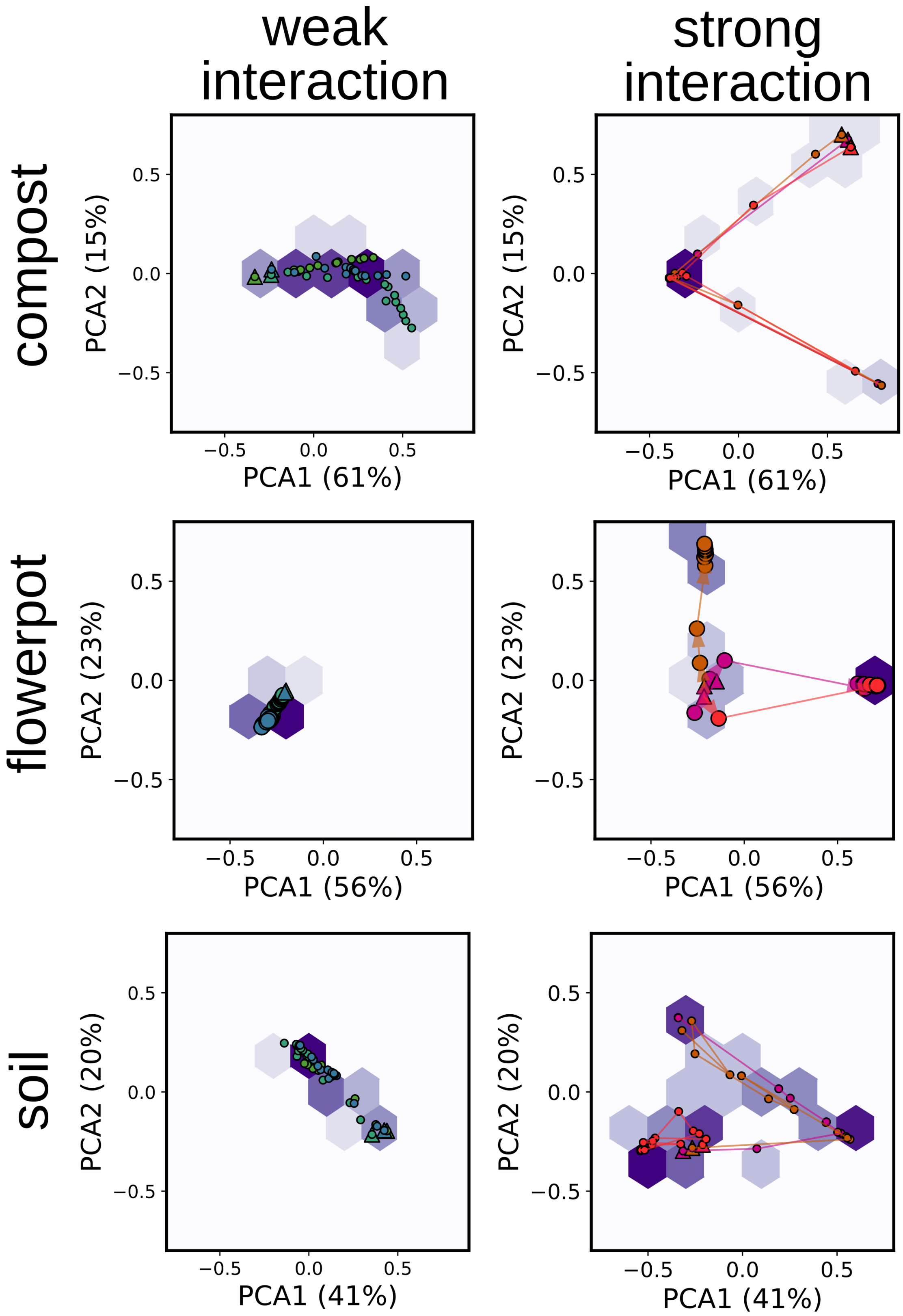


***Supplementary Figure 15: PCAs for all obtained time courses.*** *Different color shades show different replicates. Triangles show position on first day. Lines or arrows are connecting the datapoints in order of progressing time. Markers show all compositions over time. Purple hexbins corresponds to histograms with a logarithmic scaling.*

**Mathematical model of species interaction**

To describe the dynamics of the microbial communities a differential equation model was set up. This model describes the growth of a set of bacteria that interact via modifying and reacting to the environment. It thus is a multispecies extension of a model we previously used to understand pairwise interactions[1]. Further differences are that in the model shown here a heaviside function is used to describe the dependence of growth on the environment, the bacteria are not bound in how far they can change the environmental parameter p and there is periodic dilution in the system. In this model the bacteria grow with logistic growth, but just within a certain window of the environmental parameter p. If the values of p become to unsuitable for the bacteria they start to die. On the other hand the bacteria influence the environmental parameter p.

The model describes the dynamics of the bacterial species $\vec{n}'=n'_1,n'_2,n'_3,\ldots,n'_k$ and the environmental parameter p according to the formulas:

$$\frac{\partial n'_i}{\partial t}=n'_i\left(1-\frac{n'_i}{K}\right)\left[\Theta\left(-|p-p_{o_i}|+p_c\right)\left(k_{growth}+k_{death}\right)-k_{death}\right]-\delta\left(\sin\left(\pi\frac{t}{\tau}\right)\right)r_{dilution}n'_i \quad (1)$$

$$\frac{\partial p}{\partial t}=\sum_i \epsilon_i' n'_i+\delta\left(\sin\left(\pi\frac{t}{\tau}\right)\right)r_{dilution}\left(\acute{p}-p\right) \quad (2)$$

, with $k_{growth}$ the growth rate and $k_{death}$ the death rate of the bacteria, Θ is the heaviside function, that becomes 1 for $p \in [p_o - p_c, p_o + p_c]$ and 0 otherwise. If this heaviside function becomes 1 the bacteria grow with $k_{growth}$, if it becomes 0 the bacteria die with $k_{death}$. The growth and death rates are kept the same for all bacterial species (for choice of parameters see below). The heaviside function is a rather strong simplification of the growth vs p behavior, but it seems a reasonable simplification given the real shape of those curves (Supplementary Fig. 1) and the exact shape of this function does not qualitatively affect the obtained results (Supplementary Fig. 17 and [1]). δ is the Kronecker delta that becomes 1 if t is an integer multiple of τ. This term describes periodic dilution of the sample with the dilution factor 1-$r_{dilution}$. $\varepsilon_i$ is taken from the uniform interval [-$c_p$, $c_p$] and describes how the bacteria change the environmental parameter p. Accordingly $c_p$ is the maximal rate by which the bacteria can change the environment. The second term again corresponds to periodic dilution, removal of medium with parameter p and replacement with p', whereas p' is equal to the start value of p. Assuming that the carrying capacity K is similar for all species (Supplementary Fig. 16 A) we can re-scale the formulas (1) and (2) with $n_i = n'_i / K$ and $\epsilon_i = K \epsilon'_i$ :

$$\frac{\partial n_i}{\partial t} = n_i(1-n_i)[\Theta(-|p-p_{o_i}|+p_c)(k_{growth}+k_{death})-k_{death}] - \delta(\sin(\pi \frac{t}{\tau})) r_{dilution} n_i \quad (3)$$

$$\frac{\partial p}{\partial t} = \sum_i \epsilon_i n_i + \delta(\sin(\pi \frac{t}{\tau})) r_{dilution}(\acute{p} - p) \quad (4)$$

We used this re-normalized formulas for the simulation. The shift of p corresponds to the shift of the proton concentration in the experiments which itself depends on the nutrient concentrations (Fig.1). The higher the nutrient concentration the stronger the pH shift. Thus we assume that varying $c_p$ in the simulation corresponds to varying the nutrient concentration in the experiment. The main text shows simplified versions of the above equations with that does not contain the daily dilution term and shows the heaviside function as a piece-wise function.

The above equations were solved with odeint package of SciPy in python[2].

**Choice of parameters:**

The parameters for equations (3) and (4) were chosen to represent the experiments as good as possible. The experiments were diluted once a day with a factor of 1/10x therefore, $\tau$ is set to 1 (day) and $r_{dilution}$ to 0.9 .

To get an estimate for the growth rates we grew the 8 interaction partners in Base, 10mM $PO_4$ and Base, 100mM $PO_4$, 1% Glucose, 0.8% Urea for 24 hours (Supplementary Fig. 16 B) and extracted the maximal per capita growth rates from the obtained data (Supplementary Fig. 16 C). The media conditions were chosen to minimize self harming effect by pH changes. We find rather similar growth rates for most strains with a mean of the growth rates of around 6/d at low and 10/d at high nutrient concentrations and thus both are in the same order of magnitude. We use 10/d for $k_{growth}$ for the simulation.

In our simplified model we have a fixed death rate $k_{death}$, whereas in reality the death rate likely depends on the the environmental parameter in a more sophisticated way. However, to get an impression of such a death rate we look at the data published in [3] which measured the self-inflicted death by pH of bacteria and find that the death rate is similar to the growth rate. Therefore we use a $k_{death}$ of 10/d in the simulation.

The initial p and p' (the p of the replenished media) is set to 7. $p_{oi}$ is chosen from a uniform distribution in the interval [2.5, 9.5]. $p_c$ was set to 2.5. The initial bacterial density $n_i$ was set to 0.01 for each species unless stated otherwise, which corresponds to a 1/100x dilution from a saturated solution just like in the experiments. $c_p$ sets the maximal strength the bacteria change the environment and thus the interaction strength. It is varied in the simulations.

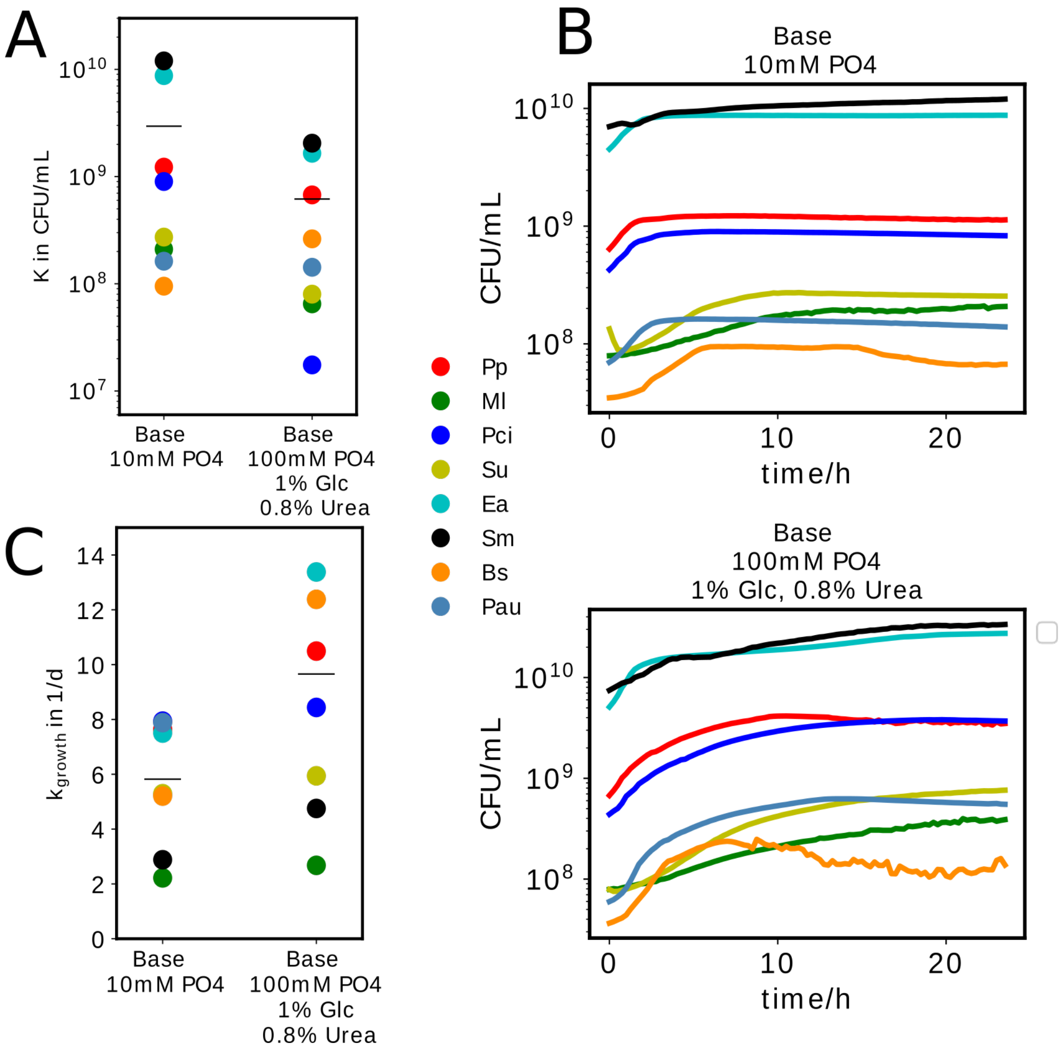


*Supplementary Figure 16:* ***Measurements to obtain simulation parameters.*** *(A) Carrying capacity for the 8 interaction strains in low and buffered high nutrient media. The conditions were chosen to minimize self inhibition of the bacteria by pH. However, at high nutrients at seems that the buffer could not fully stop self-harming pH changes. (B) Growth of the bacteria in low and buffered high nutrient media. The lines are the means of 6 technical replicates. The CFU/mL were calculated from OD with a conversion factor obtained from (A). (C) From those growth curves the maximal growth rates could be obtained. The horizontal lines in (A) and (C) show the means over all species. The single points in (A) and (C) are the means of four technical replicates.*

### Effect of interaction strength on diversity

To model how interaction strength impacts coexistence and biodiversity we numerically solved the above equations (3) and (4) for pairs and communities with 20 members. $c_p$ was varied to change the interaction strength. To make the outcomes comparable to the measurements the diversity

$^1D = \exp(-\sum_{i=1}^{S} p_i \ln p_i)$ was calculated. In the case of all species going extinct $^1$D was set to 0. The simulation were run for 80 days - which is longer then the experiments – to exclude possible 'transient' effects at early simulation times.

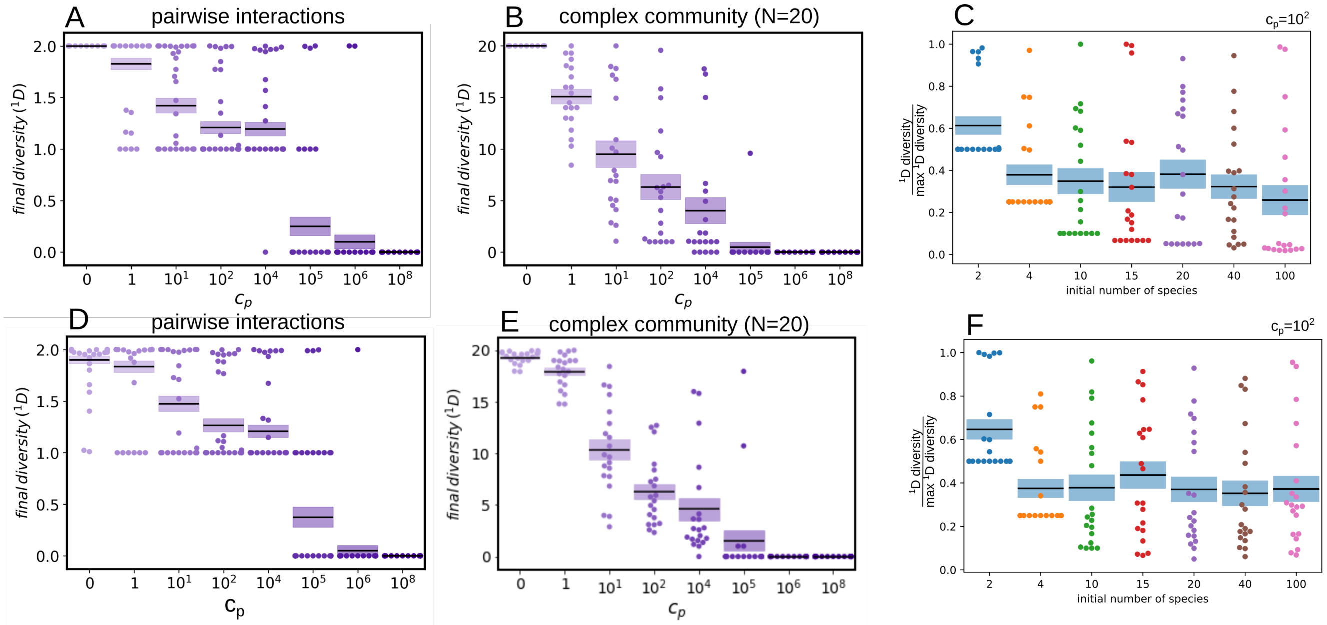


***Supplementary Figure 17: Coexistence decreases with interaction strength in pairwise interactions (A) as well as complex communities (number of initial species = 20) (B)***. *Diversity is plotted after 80 simulated days. For every interaction strength* $c_p$ *40 simulation runs for the pairs and 20 runs for the complex communities with randomly chosen* $\varepsilon_i$ *and* $p_{oi}$ *were performed as described above. The increase in* $c_p$ *corresponds to an increase of the environmental change by the bacteria and thus the interaction strength. In the shown swarm plots sometimes single data points overlap, therefore there are not necessarily 40 dots visible for every condition. (C) Relative final diversity is largely independent of initial number of species. Final diversity normalized by the maximal possible diversity (= initial number of species) is shown after 80 simulation days, 20 replicates for each initial number of species. After a slight initial drop the normalized diversity stays roughly constant with the number of species in the simulated community. However, the initial drop is likely caused by the fact, that the diversity how we defined it here cannot reach values between 0 and 1 or 0 and 1/number of species for the normalized diversity. (D), (E) and (F) are analogues to the figures above, but use a gaussian function instead of the heaviside function to describe how the bacterial growth depends on the environment.* $p_c$ *becomes the sigma of the gaussian in this case. As can be seen the results are very similar, which shows that they do not depend on the exact choice of this function.*

### Ecosystem stability

#### Stability of total biomass and environment

To estimate the effect of environmental modification and interaction strength on the total biomass ( $\sum_{i=1}^{N} n_i$ ) and the environmental parameter p over time the above differential equations (3) and (4) were integrated for 80 days with varying values of $c_p$. For every $c_p$ there were 20 different interaction pairs simulated, eg the simulation was repeated 20 times with different randomly chosen values for $\varepsilon_i$ and $p_{oi}$. The obtained results in Figure 4 and Supplementary Fig. 11 show that an increase in environmental modification leads to a decrease in stability of the total biomass.

**Stability of total biomass and environment p for pairs**

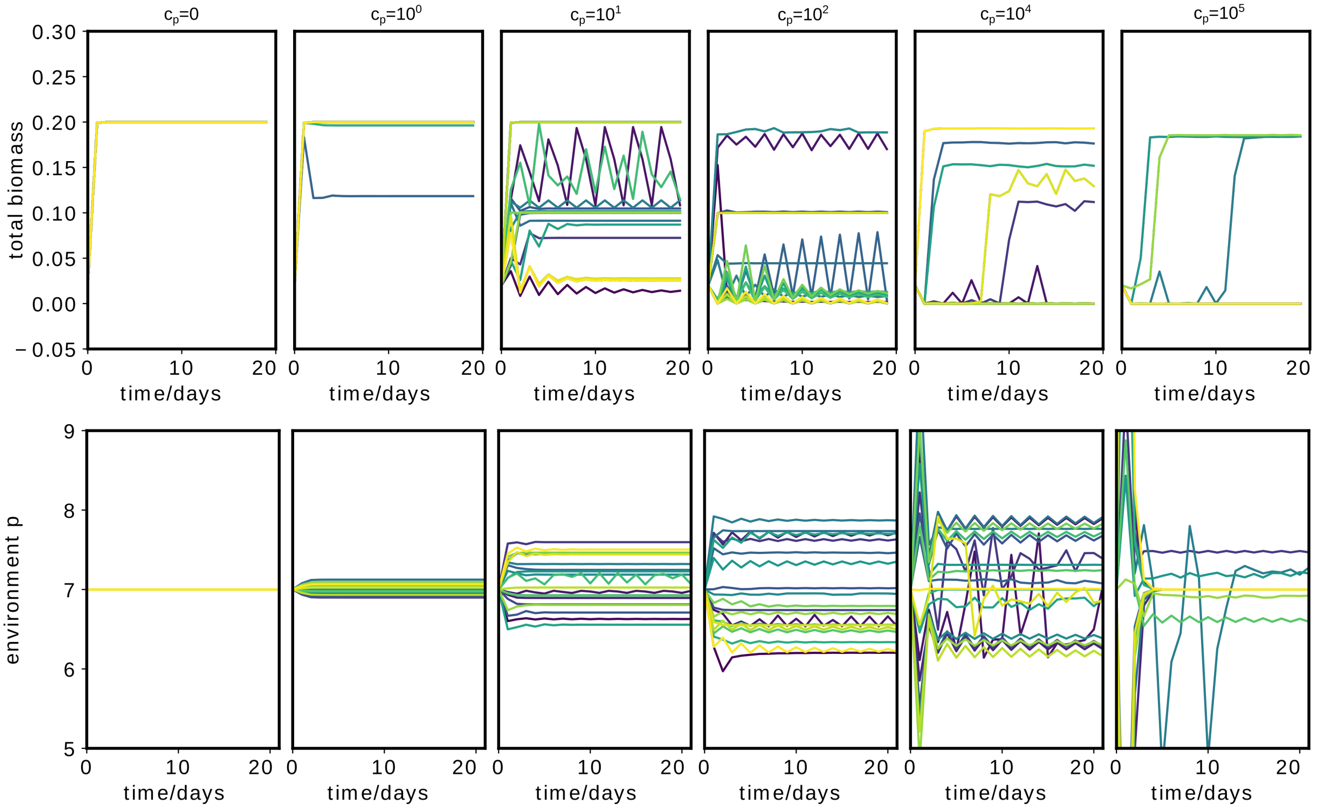


***Supplementary Figure 18: Stability of total biomass and environment decreases with increasing interaction strength in pairwise interactions.*** *Every curve (20 for each plot) corresponds to a simulation with randomly chosen values for $\varepsilon_i$ and $p_{oi}$ as described above. The dynamics of both the total biomass as well as the environment increase with interaction strength. For very strong interactions the curves show less dynamics since the bacteria often die out completely. The mean total bacterial density in the simulation is lower for strong then weak interactions, which cannot be found in the measurement (Fig. 4). However, one has to take into account that the OD measures all bacterial cells (live and dead) whereas the simulation just shows living cells. The CFU which measures only the living bacteria are indeed lower for strong then weak interaction strength as the model suggests (Supplementary Fig. 16A)*

### Stability of total biomass and environment p for complex system

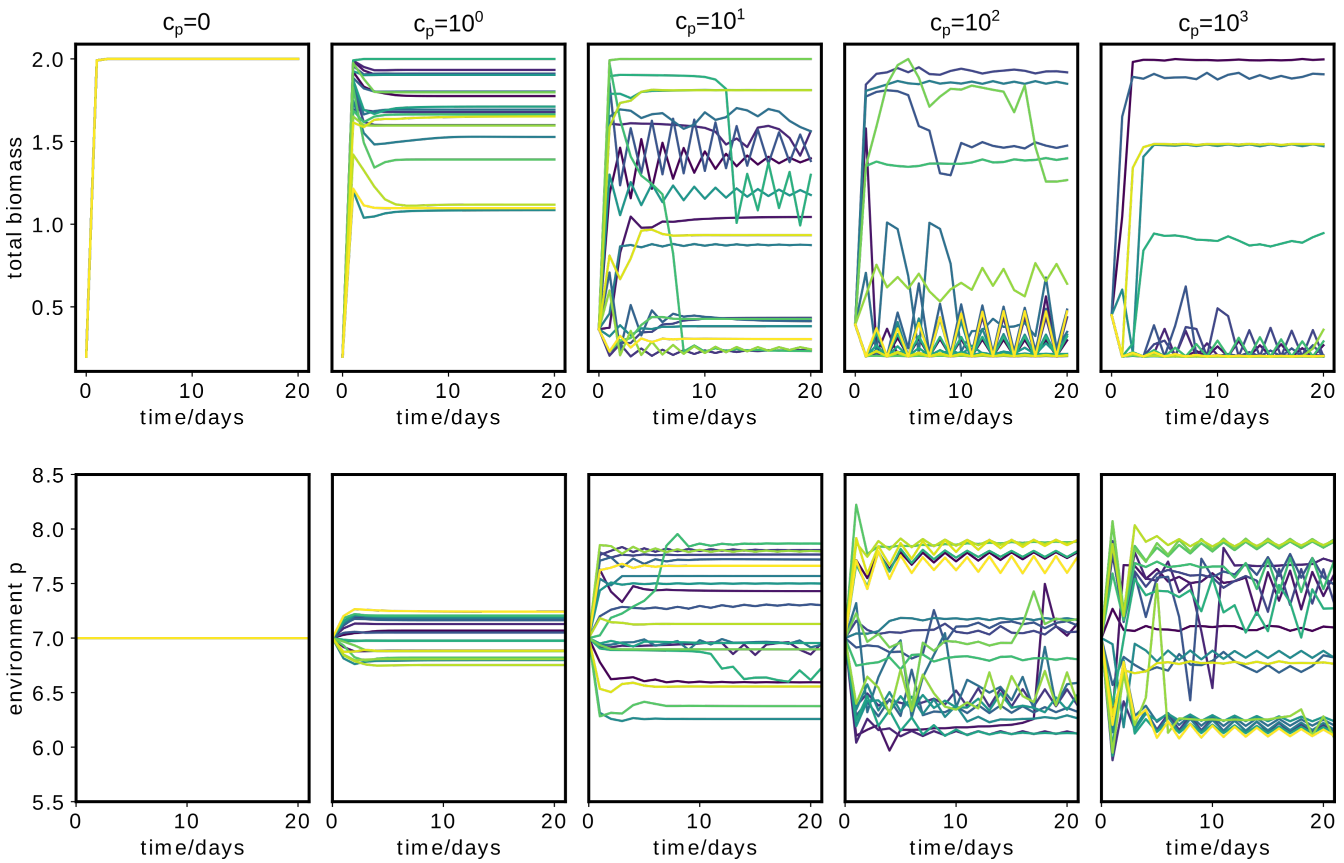


***Supplementary Figure 19 : Stability of total biomass and environment decreases with increasing interaction strength in complex communities.*** *Every curve (20 for each plot) corresponds to a simulation of 20 interacting species with randomly chosen values for $\varepsilon_i$ and $p_{oi}$ as described above.*

### Stability of species composition for different interaction strengths

### Stability of species composition for pairwise interaction

To estimate the effect of environmental modification and interaction strength on the species composition of pairwise interactions over time the above differential equations were integrated for 10 days. $c_p$ was chosen to be $10^0$ or $10^2$ for weak and strong interactions respectively. For every $c_p$ there were 90 different interaction pairs simulated, eg the simulation was repeated 90 times with different randomly chosen values for $\varepsilon_i$ and $p_{oi}$. For every interaction pair the simulation was started with different initial population densities of the interaction partners. 30 of those runs are shown in Supplementary Fig. 20 for weak interaction and Supplementary Fig. 21 for strong interactions. A summary of the outcomes for pairwise interaction analoguous to Fig. 4 in the main text can be seen in Supplementary Fig. 11.

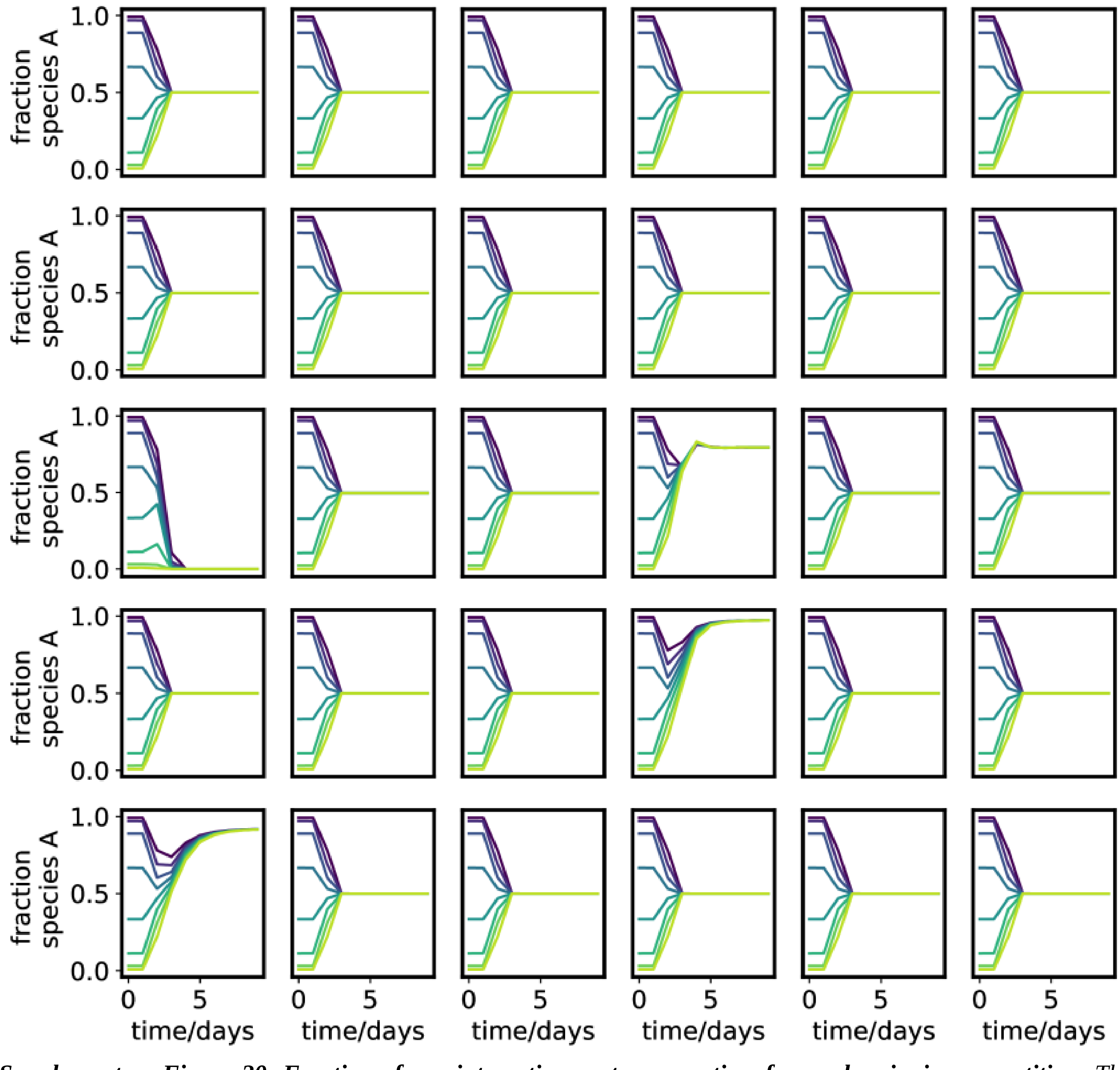


***Supplementary Figure 20: Fraction of one interaction partner over time for weak pairwise competition.*** *The simulation was started with different initial ratios of the two interacting species, each plot corresponds to a different interaction pair, eg* $\varepsilon_i$ *and* $p_{oi}$ *were chosen randomly. However, at low interaction strength (*$c_p=10^0$*) the system usually converges to one stable fixed point.*

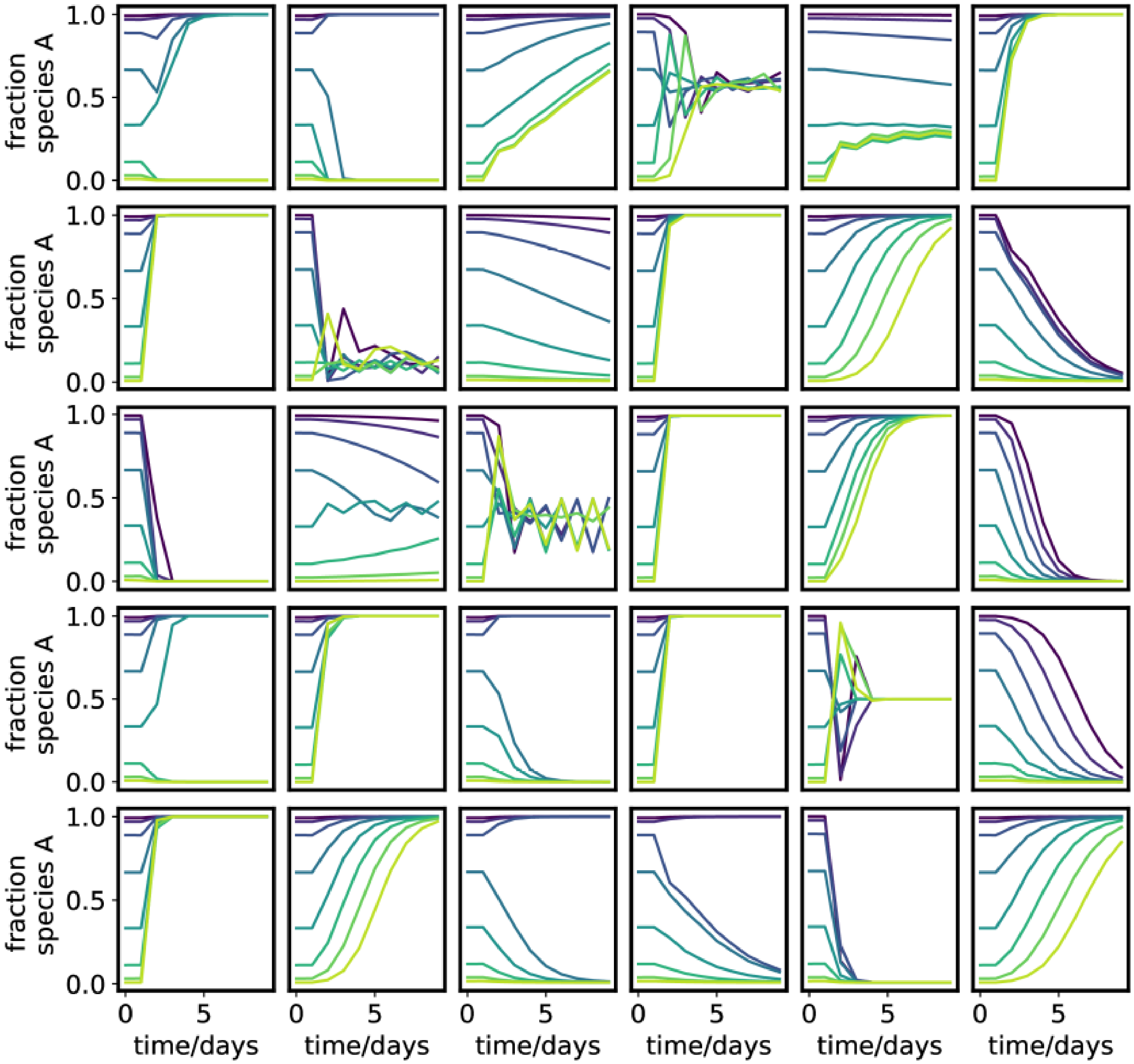


***Supplementary Figure 21: Fraction of one interaction partner over time for strong pairwise competition.*** *The simulation was started with different initial ratios of the two interacting species, each plot corresponds to a different interaction pair, eg* $\varepsilon_i$ *and* $p_{oi}$ *were chosen randomly. At high interaction strength (*$c_p=10^2$*) the system shows often fluctuations or bi-stable outcomes.*

**Stability of species composition for a 20 member community**

To estimate the effect of environmental modification and interaction strength on the species composition of a 20 member community over time the above differential equations were integrated for 80 days. $c_p$ was chosen to be $10^0$ or $10^3$ for weak and strong interactions respectively. There were 10 'systems' simulated, whereas for every system a set of $\varepsilon_i$ and $p_{oi}$ was randomly chosen. For every system the simulation was run 50 times for the both values of $c_p$ with different initial bacterial densities $n_i$ for every repeated run. The initial $n_i$ were chosen from a uniform distribution in the interval ]0,0.01]. For visualization we did a principal component analysis of the data. All obtained data-points over time for a given 'system' are shown in Supplementary Fig. 22 and one example in Fig. 4. The color shading of the hexagons shows the logarithm of the relative occurrence of the system in that area. For all subsequent compositions over time (composition at a given day and the following day) the Pearson's correlation coefficient was calculated for weak and strong interactions and plotted in Fig. 4C top, right.

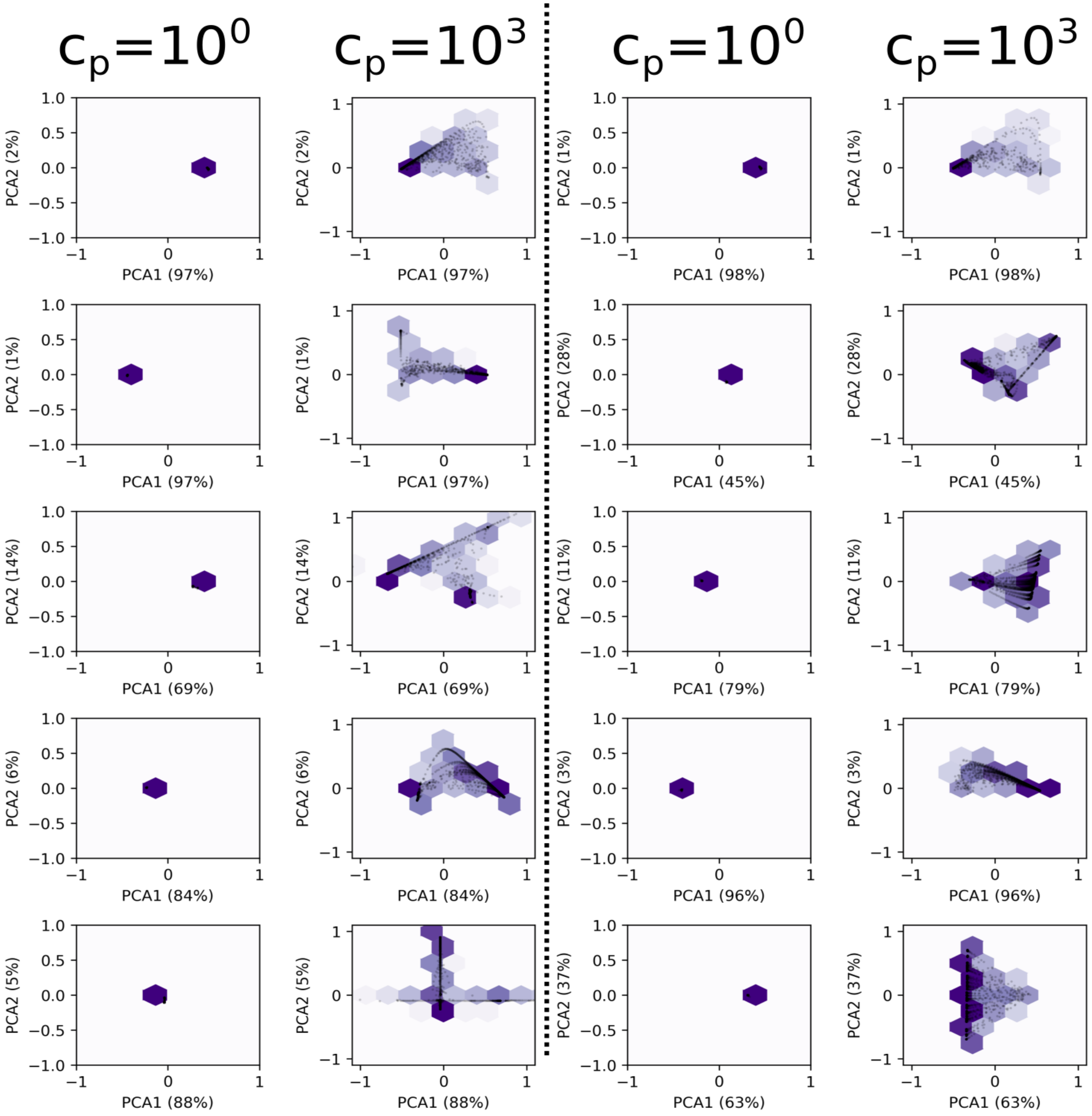


***Supplementary Figure 22: Temporal change of composition in 10 different 20 member communities for weak ($c_P$=10$^0$) and strong interaction ($c_P$=10$^3$).*** *Each system was simulated 50 times for each interaction strength. At low interactions the systems rapidly converge to a single state, whereas at higher interaction strength the systems fluctuate more strongly and show in many cases multistability. Black points show all compositions over time. Purple hexbins corresponds to histograms of black dots, with a logarithmic scaling.*
